# A functional network model for body column neural connectivity in *Hydra*

**DOI:** 10.1101/2024.06.25.600563

**Authors:** Wilhelm Braun, Sebastian Jenderny, Christoph Giez, Dijana Pavleska, Alexander Klimovich, Thomas C. G. Bosch, Karlheinz Ochs, Philipp Hövel, Claus C. Hilgetag

**Affiliations:** 1nstitute of Computational Neuroscience, Center for Experimental Medicine, University Medical Center Hamburg-Eppendorf, Hamburg, Martinistraße 52, 20246 Hamburg, Germany; Chair of Digital Communication Systems, Ruhr-Universität Bochum, Universitätsstraße 150, 44801, Bochum, North Rhine-Westphalia, Germany; Zoological Institute, University of Kiel, Christian-Albrechts-Platz 4, 24118 Kiel, Germany; The Francis Crick Institute, London NW1 1BF, UK; Theoretical Physics and Center for Biophysics, Saarland University, Campus E2 6, Saarbrücken, 66123, Germany; Department of Health Sciences, Boston University, 635 Commonwealth Avenue, Boston, Massachusetts 02215, USA

## Abstract

*Hydra* is a non-senescent animal with a relatively small number of cell types and overall low structural complexity, but a surprisingly rich behavioral repertoire. The main drivers of *Hydra* ’s behavior are neurons that are arranged in two nerve nets comprising several distinct neuronal populations. Among these populations is the ectodermal nerve net N3 which is located throughout the animal. It has been shown that N3 is necessary and sufficient for the complex behavior of somersaulting and is also involved in *Hydra* feeding behavior. Despite being a behavioral jack-of-all-trades, there is insufficient knowledge on the coupling structure of neurons in N3, its connectome, and its role in activity propagation and function. We construct a model connectome for the part of N3 located on the body column. Using experimental data on the placement of neuronal somata and the spatial dimensions of the body column, we show that a generative network model combining non-random placement of neuronal somata and the preferred orientation of primary neurites yields good agreement with experimentally observed distributions of connection distances, connection angles, and the number of primary neurites per neuron. Having validated the N3 connectome model in this fashion, we place a simple excitable dynamical model on each node of the body column network and show that it generates directed, short-lived, fast propagating patterns of activity. In addition, by slightly changing the parameters of the dynamical model, the same structural network can also generate persistent activity. Finally, we use a neuromorphic circuit based on the Morris-Lecar model to show that the same structural connectome can, in addition to through-conductance with biologically plausible time scales, also host a dynamical pattern related to the complex behavioral pattern of somersaulting. We speculate that such different dynamical regimes act as dynamical substrates for the different functional roles of N3, allowing *Hydra* to exhibit behavioral complexity with a relatively simple nervous system that does not possess modules or hubs.

## 1 Introduction

### 1.1 Information processing in neuronal systems

Information processing in biological neural systems takes place on the stage of complex neuronal networks (1). These networks exhibit vast structural and functional diversity, ranging from primate brains with billions of neurons, tens of billions of synapses, and a myriad of structured and functionally specialized sub-networks to nervous systems with low structural complexity, as exemplified by the nematode *C. elegans* with conserved number, position, and function of every neuron (2).

A fundamental principle common to all nervous systems, however, is given by the notion of neural computation, during which sensory organs of an animal receive information from its surroundings. Then, the information is transmitted to a particular nervous system that integrates it and finally generates output that leads to animal behavior.

Apart from these general principles, there often is no consensus on the precise details of a neural computation. In fact, it is frequently not even clear which neurons are precisely involved, how they are connected, and how their activity contributes to a given neural computation. The main reason is that the resulting networks and their connectivity are very complex and experimentally difficult to access. This makes a systematic and mechanistic understanding of neuronal computations an arduous task.

### 1.2. *Hydra* as a model system for neuroscience

To partially circumvent these difficulties, in recent years, there has been a research effort directed at understanding presumably simpler nervous systems with a focus on aquatic invertebrates forming the phylum *Cnidaria* (5), which includes jellyfish (6–8), anemones, corals, and polyps. Among the polyps, the small freshwater cnidarian *Hydra* has received a lot of interest in neuroscience research (5). *Hydra* is a transparent animal with low cellular complexity and a simple body plan (Fig. 1 (a)): Its tube-shaped body column is made up of three cell layers: the ectoderm, mesoglea, and the endoderm. At the bottom of the animal is a peduncle, whereas at the top, hypostome and tentacles are located. This long axis of the animal is also called the oral-aboral axis or longitudinal axis. The nervous system of *Hydra* consists of two distinct nerve nets, one each in ectoderm and endoderm. Recent transcriptomic analysis showed that there are 7-11 neuronal populations in *Hydra* (3,4) (cf. Fig. 1 (b)). On a functional level, a seminal study revealed that this genetic specification is, at least in part, mirrored by the fact that *Hydra* behavior is controlled by non-overlapping, functionally distinct neuronal populations (9). Interestingly, neuronal processes in *Hydra* do not show a differentiation into axons and dendrites, like in vertebrates, but instead are morphologically unpolarized structures called neurites (9). Despite its simplicity, *Hydra* exhibits a range of neuron-dependent behaviors (10), such as feeding (11,12), elongation of its body column in response to stimulation with light and somersaulting (13). Although there is this wealth of behavioral data, there is not much known about the structural basis of *Hydra* behavior. Early work using electron microscopy (14,15) showed that there are both chemical and electrical (gap junctions) synapses in *Hydra*, but did not make an attempt to elucidate the structure of the underlying neuronal networks (16).

**Fig. 1:**
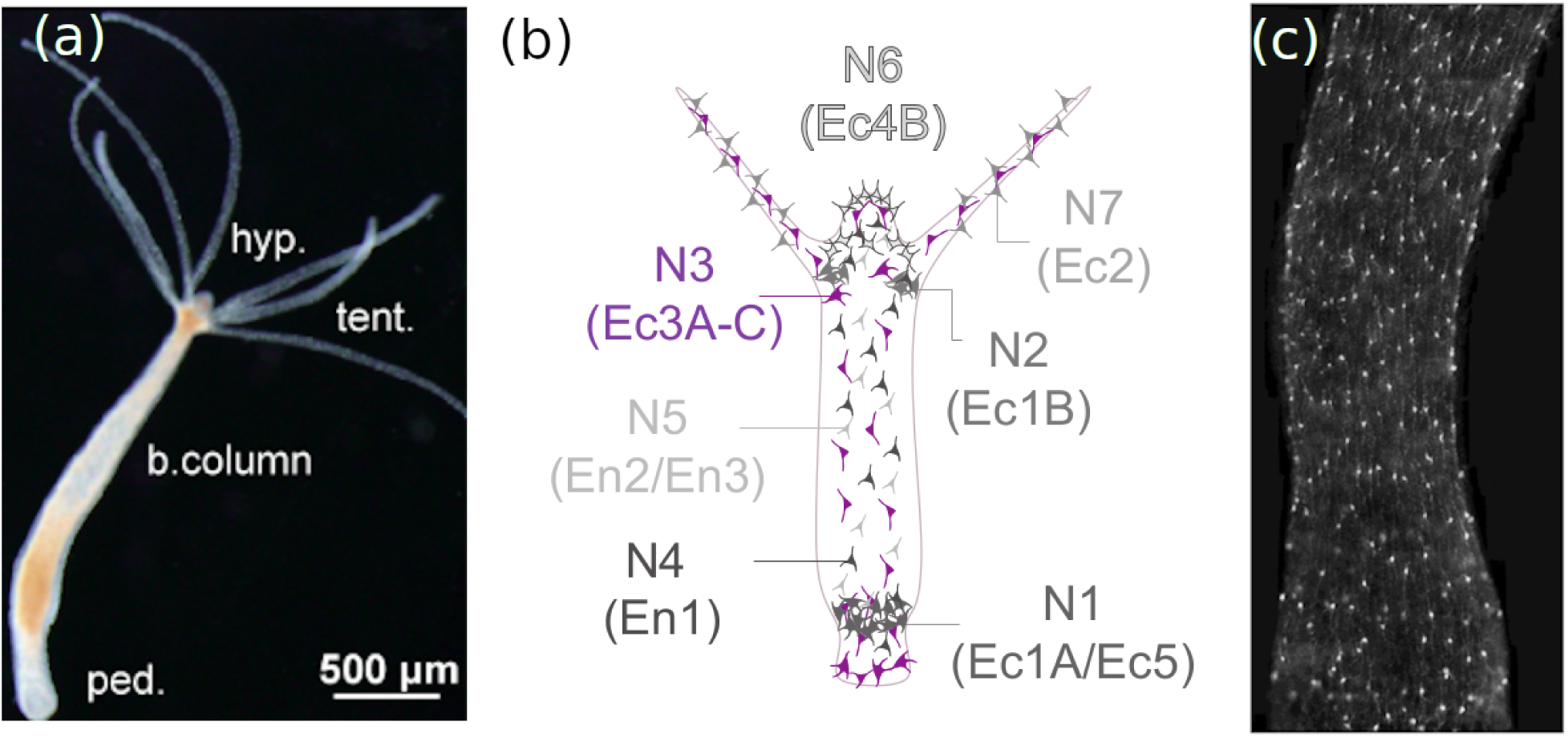
Overview of *Hydra*. (a) *Hydra* polyp in relaxed form, showing an elongated body shape with peduncle (ped.), body column (b.column), tentacles (tent.) and hypostome (hyp.). (b) Schematic presentation showing location of seven neuronal subpopulations and their distribution (after (3,4)). The alternative nomenclature includes Ec for ectoderm and En for endoderm. N3 (Ec3A-C), highlighted in magenta, is located throughout the polyp. (c) Image of population N3 located in the body column. Hypostome and peduncle are not included. Soma are visible as bright spots. Neurites (thin bright lines) extend mainly in the up-down (oral-aboral) direction.

One exciting neuronal population, population N3 (Ec3), can be found in the ectoderm of *Hydra* (4) (Fig. 1 (c)). This population was recently shown to be modulating *Hydra* feeding behavior, playing a role in internal state, phototaxis, and somersaulting (12,13). In the complex behavior somersaulting, N3 increased its activity in a ramp-like fashion, culminating in a series of population bursts which finally initiates the foot detachment and the somersault.

In light of these observations, the question arises whether and how a single neuronal population can support different modes of neuronal activity. Based on microscopy images, the organization of the nerve net N3 is not random, but follows some regularity (Fig. 1 (c)). Therefore, here we aim to determine the structural basis for the N3 subnetwork and study how this structure can give rise to different modes of activity propagation, which likely underlie different behaviors of *Hydra*.

### 1.3. Outline of the paper

This paper is structured as follows. We first construct a theoretical connectome model respecting basic geometric constraints. To achieve this, we introduce a statistical model that faithfully approximates intersomatic distances recorded in microscopy images of N3 neuronal somata. Based on a representative distribution of soma locations on a two-dimensional domain, we then construct a theoretical connectome following pairwise location- and orientation-dependent connectivity rules. We determine how the derived model compares to experimentally measured values for connection length, connection orientation, and number of connections per neuron. In the second part of the manuscript, we study neural activity on our theoretical connectome. First, we show how the connectome model combined with a simple model for excitable dynamics can give rise to two qualitatively different activity patterns. Second, we study neural activity using a version of a previously published single neuron model (17). By construction, this model allows for a neuromorphic circuit replicating the neural network dynamics. We investigate under which circumstances our theoretical nerve net gives rise to two distinct neuronal activity modes, provide a mechanistic explanation for this and discuss the biological plausibility of our findings.

Our work constitutes the first step to systematically construct and validate a connectome for *Hydra*, starting with the body column ectodermal population N3.

## 2. Results

### 2.1. Constructing a statistical model for the location of neuronal somata

To generate a statistical model for the location of neuronal somata and compare it to the biological samples, our first task is to study the location of neuronal somata from population N3 in the body column of the animal. N3 is an ectodermal population, so all its somata are placed in the ectoderm. *Hydra’s* body is a three-dimensional cylinder. This means that somata are placed on a cylinder shell. However, *Hydra* shows rotational symmetry around its long axis (from hypostome to peduncle or vice versa), and therefore, considering one side of the cylinder suffices to study the structure of N3. Imaging (see Methods) was therefore performed in one imaging plane on one side of the animal. Thus, throughout this paper, we assume that neurons are placed on a two-dimensional, quadrilateral domain, thus neglecting the fact that *Hydra’s* body column is shaped like a cylinder in three dimensions. We also assume that somata do not possess a spatial extent, that is, we consider neurons as points in a two-dimensional space.

For notational convenience, we use quadrilateral domains with side lengths Δ*x* and Δ*y* in the *x-* and *y*-direction, respectively. *Hydra* has a larger length than width, i.e., Δ*y = α*Δ*x* with *α >* 1.

Thus, the total area of the domain is *A* = Δ*x*Δ*y* and therefore, 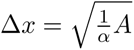 and 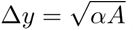. We consider *N* neurons with coordinates 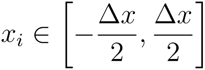 and 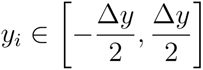, for the theoretical connectome.

Which rules could govern the placement of neurons on a two-dimensional part of *Hydra’s* body column? Phenomenologically, neurons have a certain minimal distance of around 10*μm* from each other, but otherwise are not arranged in a regular fashion. This entails that neuron positions are not random, but follow some other principles of placement. To see why positions cannot be random, assume that for each neuron, its *x* and *y* coordinates are drawn uniformly and independently (both independently from neuron to neuron as well as independently between *x* and *y* coordinates for a single neuron) from the intervals given above. This would lead, with some probability, to neurons that are placed arbitrarily close to each other, which disagrees with the experimentally observed data.

How different are the observed neuron positions from random placement? To address this question, we turn to the algorithm of Poisson disk (PD) sampling (18). In this algorithm, points are placed on a domain of a given size with the aim that no two points are closer than a distance *r*, that is, *r* defines the minimal distance between any pair of points. Distance here is defined as the standard Euclidean distance between two somata. When we naively apply this algorithm on domains that match measured ones in terms of area, we find that too many points (neurons) are needed when we work with biologically plausible minimal distances. This placement of neurons also led to a fairly regular placement of points compared to the experimental data, entailing that the typical distance between two points was given by the minimal distance. Thus, in a second attempt, we delete the number of surplus points, which preserves the observed minimal distance. However, we find that this still does not result in a better-than-random fit to the experimentally determined intersomatic distances, as quantified by the pairwise intersomatic Euclidean distances. In a third attempt, instead of specifying the minimal distance *r*, we specify the number of points that are to be placed on the domain, and then determine the corresponding minimal distance.

Which number of points should be chosen? To quantify this, we consider the distributions of pairwise intersomatic Euclidean distances, and compare distributions of the experimental data (soma locations for different polyps, see Methods for details of data acquisition) to those obtained with our PD sampling approach. We find that it must be larger than *N,* the target number of neurons determined by experimental data (see below), but not too large, so that it still results in a larger minimal intersomatic distance than in the experiment. We choose *2N* as the initial number, so that we need to remove *N* points later. This approach already yields good agreement of the model with the data. A final correction is performed after the removal of the surplus points, because we find that our algorithm up to this point produces a minimal intersomatic distance that is too large. Thus, a random subset with a fixed size of 50 neurons is chosen, and the coordinates are re-sampled randomly, but only accepted when the new coordinates were not closer to already existing points than the minimal distance. This approach yields the best agreement with the experimental data (see Fig. 2 (c) for one example of experimental data and Fig. 2 (a) and (b) for PD and random sampling, respectively), and better approximates intersomatic distances than random sampling (compare Fig. 2 (c) to 2 (a) and (b)).

**Fig. 2:**
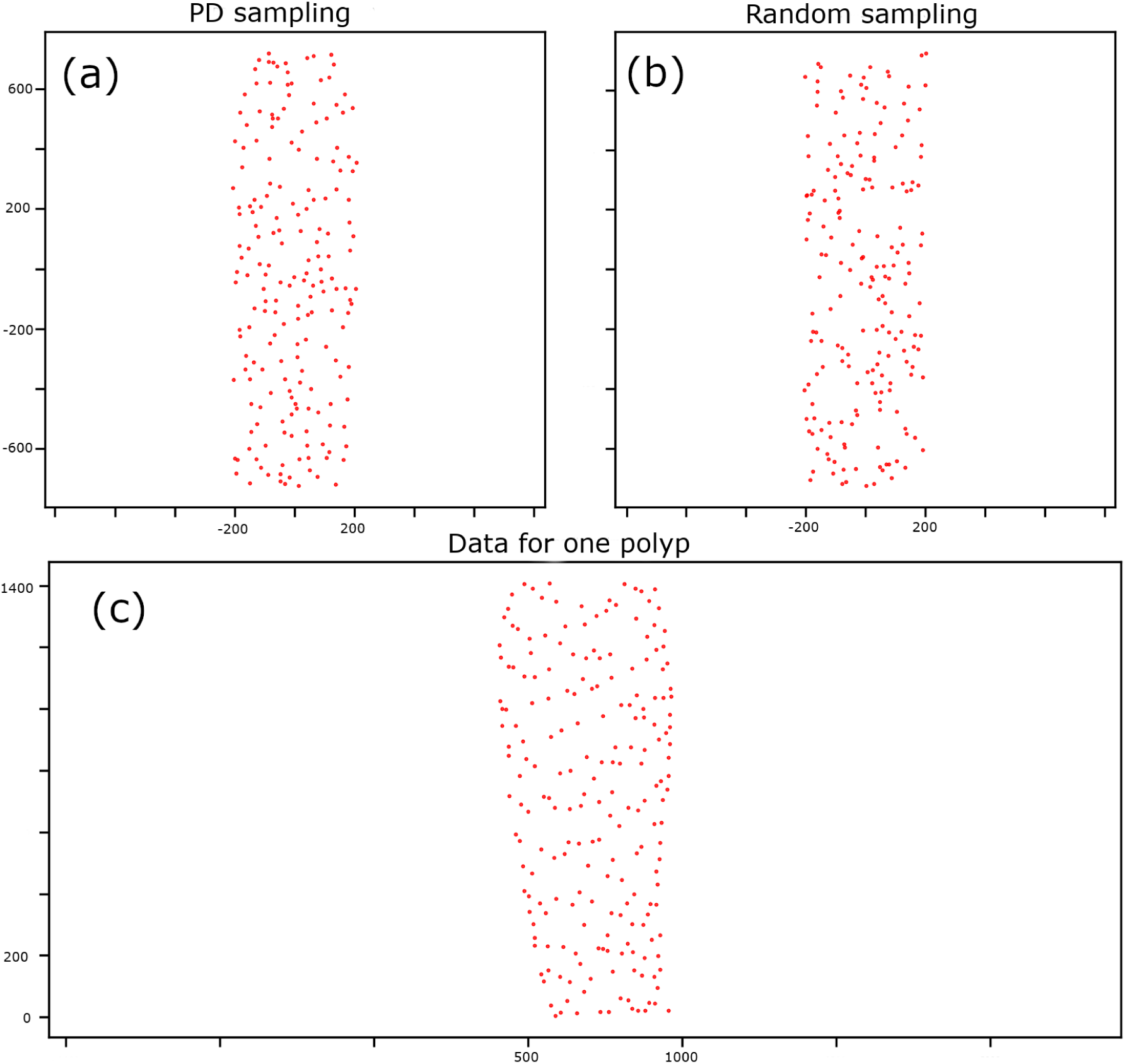
Constructing a model for the location of neuronal somata. (a) Poisson Disk (PD) sampling. (b) Random sampling. (c) Data for one polyp. Spatial dimensions are given in A^im^. Note that random sampling results in more clusters of somata that are close to each other, compared to PD sampling and also compared to the data shown in (c).

After testing this algorithm on each of the *M* = 15 measurements of neuron positions individually, we decided that it is most advantageous to construct a single representative distribution of points so that we later only have a single representative connectome. We thus average neuronal densities, areas and minimal intersomatic distance and applied our modified Poisson distance algorithm with the same parameter for each of the experimentally measured datasets. We also chose *α* = 3.5 for each data set. This results in a computational domain with a width of approximately 414 and a height of approximately 1450 *μm* (Fig. 2 (a)), which means that the area is approximately 0.6 mm^2^. The average neuron density in the dataset is 321 mm^-2^, which resulted in *N* = 192 neurons placed on the quadrilateral computational domain.

We then quantify the agreement of the distributions for pairwise intersomatic distances obtained from the model with the data (Fig. 3). We restrict the analysis to intersomatic distances up to and including 250 *μm* to increase the sample size as not all datasets had the same spatial extent. While the agreement, as quantified by large p-values from a 2-sample Kolmogorov-Smirnov test, is good across the entire dataset, and better than random, it can for some datasets yield to small p-values which indicates that the fit is not good (Fig. 3 (b), part of distribution below red dashed line). This is likely due to the fact that some of the average parameters, determined as means across the entire population, are not representative for some of the data. Indeed, there is a large degree of variability in the dataset, reflected in both the shape of the polyps as well as their dimensions and the number of neurons, which ranges from 259 to 120 in our dataset. The density, however, was more constant: for a large animal with an area of 0.9 mm^2^ and 259 neurons, it is approximately 287 mm ^2^, whereas for a smaller animal with only 120 neurons and an area of 0.4 mm^2^, it is approximately 299 mm^-2^. Therefore, our approach of focusing on neuron densities takes into account the large variability in the dataset. In each case, running our algorithm with mean parameters gives good agreement of the minimal intersomatic distance and always reasonably approximates all datasets (Fig. 3 (b)). Therefore, we continue to work with this single distribution of points, which is fixed up to the random seed of our modified Poisson disk sampling algorithm.

**Fig. 3:**
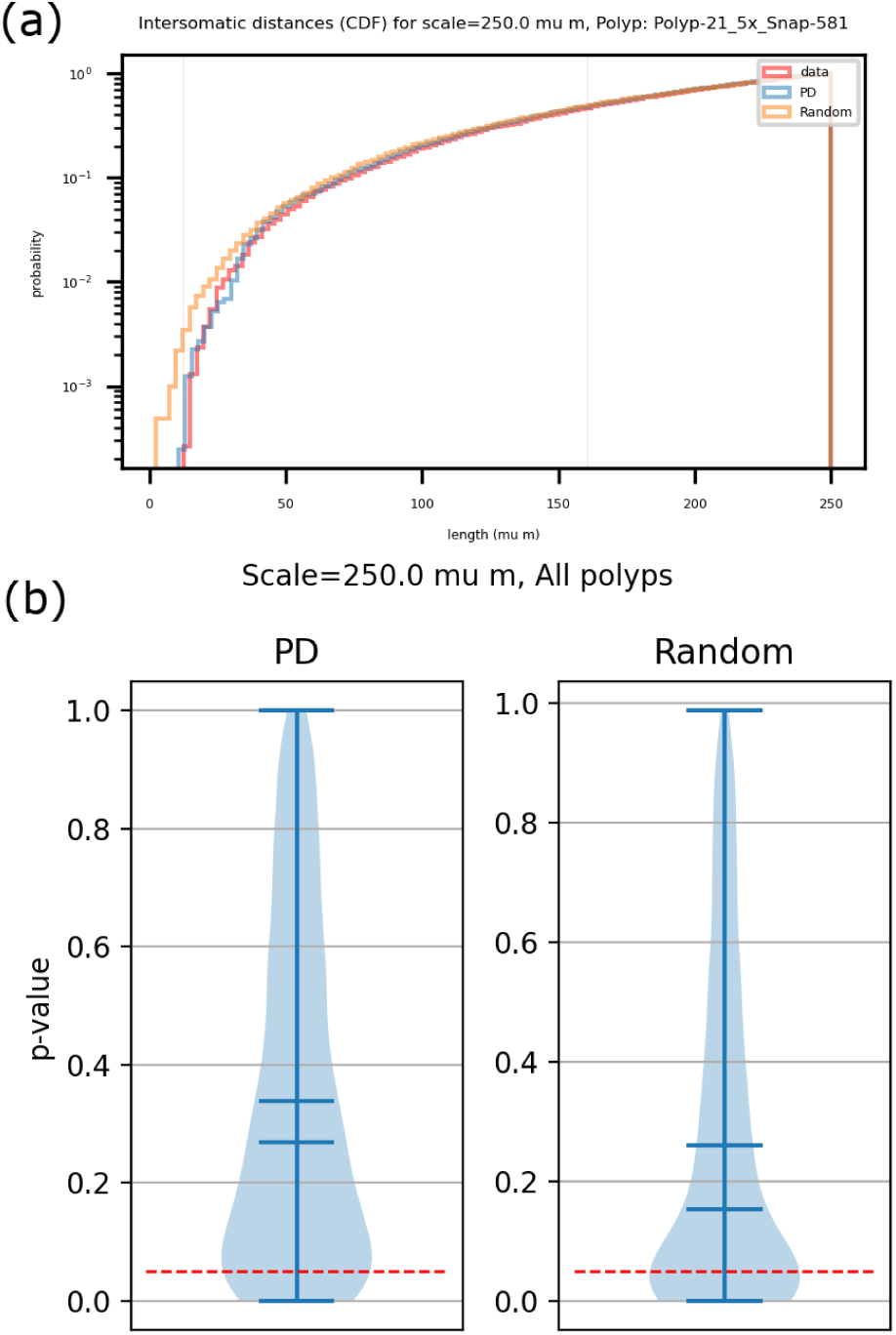
Statistical Comparison of PD and Random Sampling. (a) Cumulative distribution function (CDF) for the soma positions shown in Fig. 2 with a cut-off at 250 *μm*. Note that pd samping results in a better fit to the data distribution than random sampling, which results in a smaller minimal intersomatic distance and an overall overestimation of small intersomatic distances. (b) Distribution of p-values for two-sided two-sample KS test. Results are pooled across the whole dataset *(M* = 15 polyps) for 100 independent random realizations per polyp. The difference between PD and random sampling is significant (*p* < 10^-10^, one-sided Mann-Whitney U-teston the two shown distributions). Red dashed line indicates *p* = 0.05. The two middle horizontal bars indicate the mean and the median of the two distributions, which are both higher for PD sampling compared to random sampling.

To sum this up, our modified PD sampling algorithm takes as input the average density of neurons, the average minimal distance, the chosen scale parameter *a* and the average area of the polyps and returns the coordinates of *N* neurons on a fixed quadrilateral domain. The averages for the input quantities are taken across *M* = 15 different polyps.

### 2.2. Constructing a connectome for N3

Having defined a statistical method to place neuronal somata on a computational domain in the shape of a rectangle with biophysically plausible dimensions, we now describe our algorithm to generate an artificial connectome based on location- and angle-dependent pairwise connection rules.

A striking feature of the data is the fact that neurite orientation is not random ( however see (9) where no preferential or specific orientation was found for any the networks studied, see also Discussion), but that there is a preference for neurites to run longitudinally along the main axis of the animal, that is, from “up” (starting from below the hypostome) to “down” (towards the peduncle) or vice versa. Thus, the distribution of neurite orientation has a peak at 90°. Whether this preferred orientation is the result of spatiotemporal ontogenetic outgrowth rules (19) or constrained by the longitudinal orientation of ectodermal muscles is not clear.

Going beyond this basic structural feature, in general, neurites of N3 neurons do not run straight between two neuronal somata and have a complicated branching structure (see Appendix Fig. S2 for an example). Therefore, the direct Euclidean distance between two neurons is usually shorter than a neurite connecting them. Hence, instead of modeling complex neurite topology, we decided to connect somata with straight lines. In measurements, putative connections between neurons were determined by tracing a neurite from one soma until it finishes at another soma without crossing a third soma on the way, irrespective of whether the neurite branched or not (Appendix Fig. S2). When a connection was deemed as present, the Euclidean distance between two neurons was recorded. With only antibody stainings available (see Methods “Immunohistochemistry against GFP and Imaging” for details) and without electron microscopy imaging, definite knowledge on the presence or absence of connections is lacking and our connection distance measurements between two putatively connected neurons is only a proxy for the real presence of a connection. The experimentally measured value for the connection angle is given by the signed orientation of the line directly connecting two putatively connected neurons (Appendix Fig. S1).

Therefore, our modeling approach uses several simplifying assumptions. First, as described above, neuronal somata are placed on a two-dimensional domain, whereas real *Hydra* are three-dimensional animals. Second, all connections are straight lines connecting two somata. Hence, neurites are not modeled as the complicated twisting and turning structures they are in real animals, but instead as straight lines connecting two somata. As a consequence, our model does not consider branching: all neurites are straight lines that start and finish at two somata, which themselves are modeled as points. We use ‘neurite’ and ‘connection’ as synonyms as far as the model is concerned. Moreover, we only allow one connection between two neurons.

Overall, the aim of our modeling approach is not to reproduce the morphological complexity of N3, but to build a functional minimal network model that is in agreement with measured experimental data and can at the same time show different dynamical regimes also present in the real animal.

We choose to model connections as undirected and unweighted. Therefore, we obtain a binary symmetric adjacency matrix as the model connectome. This is justified, because contacts between neurons in N3 are mainly formed by gap junctions, but chemical synapses might also be present (4). The resulting connectome therefore is a spatially embedded graph whose nodes are interpreted as neuronal somata and whose edges are interpreted as neurites.

Before connections are made in the model, each neuron is assigned a maximum number of connections from an experimentally measured distribution counting the frequency of the occurrence of a certain number of primary neurites per neuron. In our model, we take the number of neurites as equivalent to the maximal number of connections per neuron, and hence its degree. Thus, to each neuron, we randomly assign an integer number which restricts its maximal degree. The assignment is independent of neuron position and its spatial neighbors. Given that neurites in *Hydra* often show substantial overlap (16), so that whenever two neurites cross, a putative connection between two neurons can consist of more than one synaptic contact. Further, given that two neurites can cross more than once, this assignment of maximal degree via primary neuron count is another simplifying assumption.

The model does not include a temporal dimension and there is no pruning: ontogenesis is not modeled and whenever a connection is made, it remains in the network. With neuron positions given by the PD sampling algorithm described above, the algorithm to generate the connections consists of three main steps. Each step has the form of a loop through all pairs of neurons in the network, with the exception of self-connections, which are excluded from the model. In the first two loops, so-called longitudinal model neurites with orientation close to the preferred direction are made. In the third, and final, loop, lateral model neurites with no restriction on preferred direction are made. Each loop proceeds in two main steps. First, it is checked whether a candidate connection between two neurons fulfills certain loop-specific criteria. If the criteria are fulfilled, the distance between two neurons is recorded and the connection is marked as a candidate connection. Second, after running through all potential connection partners for one given neuron, connections are made in ascending order of the distance between candidate pairs, while still checking for the connectivity criteria in each case. The criteria slightly differ between the three loops. The common criteria in each loop is that no neuron is allowed to make more connections than it was assigned from the distribution specifying its maximal degree (see above). In the first loop, the aim is to make at most two connections per neuron, of which one should point ‘upwards’ (quadrants I and II of a cartesian coordinate system with the first neuron at the center), and the other one should point ‘downwards’ (quadrants III and IV of the same coordinate system). The connection orientation for the second neuron in the loop is likewise checked in a cartesian coordinate system with its positions at the center. Connection angles are computed from the coordinates of the two neurons by using the slope of the line connecting the two neurons. The angle obtained from the slope is transformed according to the convention shown in Appendix Fig. S1 to facilitate comparison with experimental data. The orientation selectivity is implemented so that an orientation angle (see Appendix Fig. S1 for angle sign conventions) assigned to each connection is not allowed to deviate more than Δ*β* = 15° from preferred orientation 90°. For simplicity, unlike in the experimental data, we do not set different preferred orientations for neurites pointing upward or downwards, so that we do not have a clear distinction between a region close to the hypostome or a region close to the peduncle in our model. Overall, we found that this choice of Δ*β* led to a good overlap between the connection distance-angle orientation correlation when comparing data and model (Fig. 4, angle-distance correlation). Thus, after the first and second loop, connection angle orientations are restricted to ±90° ± 15° (large peaks around ±90°in Fig. 4, connection angles). Moreover, the maximal length of a connection is restricted to 450 *μm* After the completion of this loop, each neuron has at most two connections (neurites) in opposite directions which preferentially run along the long axis of the computational domain. The second loop is very similar to the first loop, with the exception that each neuron now can make at most three connections in total (counting the connections made in the previous loop as well). Moreover, not all three connections are allowed to be made in the same direction: a neuron is allowed to have two neurites pointing upwards, and one pointing downwards, but not three (or more) pointing downwards. It is also allowed to have two neurites pointing in the same direction. We allow for two neurites to be made in the same direction because of neurons located close to the border of the computational domain, which would otherwise not find connection partners. After the completion of this second loop, each neuron has at most three connections. The network with only longitudinal model neurites is shown in Fig. 5, right panel. The third and final loop places no restriction on the orientation of a neurite. The maximal connection length is now reduced to 200 *μm* and no neuron is allowed to make all connections in one direction when it has more than two connections. This last loop has two purposes: first, because even if the experimentally obtained distribution of connection orientations is peaked, not all connection angles are in a narrow interval around the preferred orientation (Fig. 4, connection angles). Instead, nearly all values of the orientation are present. Because a strict restriction on the deviation from the preferred angle was placed in the first and second loop, this would otherwise not have been reflected in the model. Second, some neurons make more than three connections, which would be impossible after the execution of only the first two loops. Indeed, whereas most neurons make 3 connections, some neurons can make up to 6 connections (Fig. 4, degree/ primary neurites) The subnetwork consisting solely of lateral model neurites is shown in Fig. 5, middle panel. The complete model connectome is shown in Fig. 5, left panel. The agreement of model and data quantities is shown in Fig. 4. We note that the output of the connectome model is fixed when the neuron positions and the maximal number of connections per neuron as well as the order in which the nodes are traversed, are given. Apart from the order in which the neurons are traversed in the outer loops, which we have fixed here, the generative algorithm as such is purely deterministic and does not contain any random elements. After the execution of these three loops, it can still be the case that a neuron only has one connection, which is the case for a small number of neurons (Fig. 4, degree/ primary neurites). We do not manually add more connections to a neuron in this case. In general, using different random seeds, it can also be that the resulting network has more than one connected component. This, however, is not the case for the specific network realization we consider in this paper (Fig. 4 and 5).

**Fig. 4:**
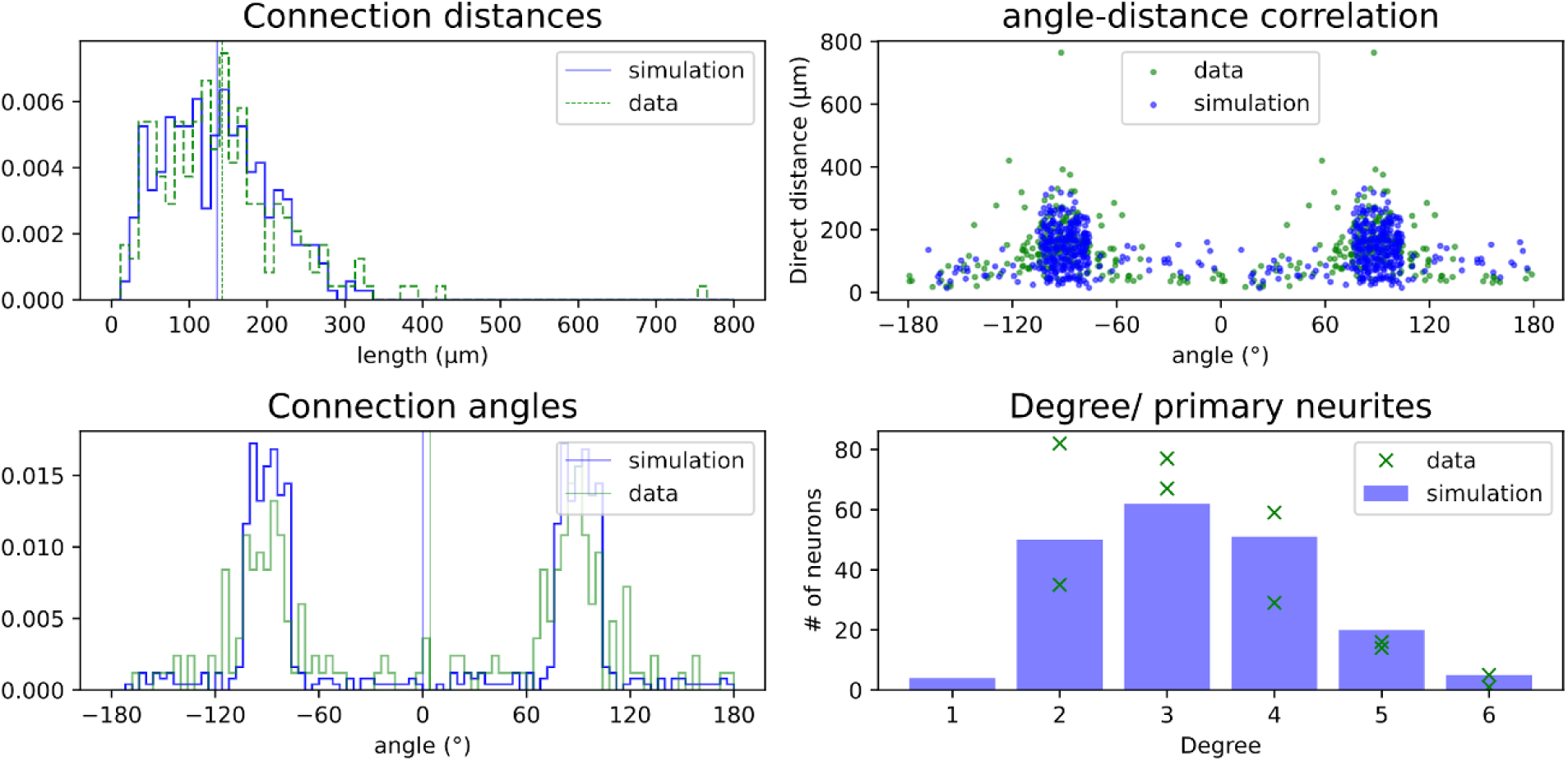
N3 connectome: Comparison to empirical data. Comparison of model quantities with data. Green: data. Blue: quantities derived from connectome model simulation. Clockwise, starting top left: Connection (direct) distances (histogram), angle-direct distance correlation, connection angles (histogram) and degree as measured by the number of primary neurites. For the degree distributions, two independent datasets were generated, and therefore, for each degree value, there are two markers, one for each dataset. The spatially embedded network model (connectome) is shown in Fig. 5.

**Fig. 5:**
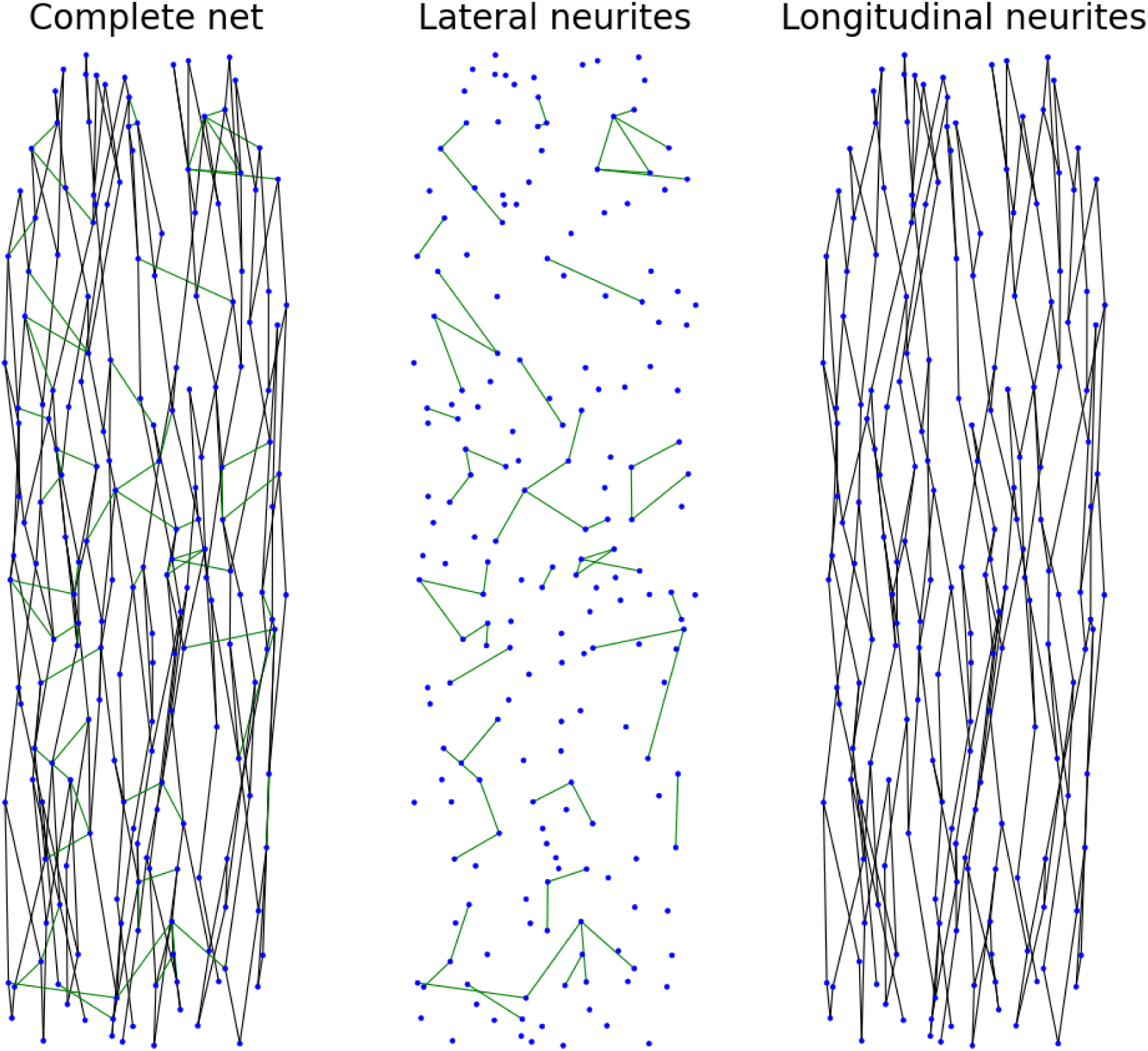
Network composition. The complete network (left) is composed of model lateral neurites (middle) and model longitudinal neurites (right), which are added to the network in three steps (see main text). Neuronal soma positions are given as blue dots. The network has 312 connections.

Our resulting network is sparse: with only *C* = 312 connections, the sparseness defined by 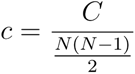, as appropriate for undirected graphs without self-connections, is approximately *c* = 1.7%. Allowing for a higher number of maximal connections would increase this value, result in more connections and hence less sparse networks.

In summary, our model connectome is an effective structural synaptic connectome. It serves as a first step to construct more detailed, and potentially more biologically plausible, models for the *Hydra* nervous system. Going beyond modeling single neuronal populations in *Hydra* is required for a better understanding of the functioning of its nervous system. Eventually, for a complete understanding of neural computations in *Hydra*, all neuronal populations must be modeled similar to N3 here, and coupled with each other, and the external environment, to be able to achieve a comprehensive biomechanical understanding of *Hydra* behavior (20).

### 2.3. Studying activity propagation with a simple minimal excitable model

With the connectome model at hand, we study activity propagation in this network, using a simple model for excitable systems which evolves in discrete time, the SER model. In this model, each node can be in one of the three states: susceptible (S), excited (E) and refractory (R). Whenever a susceptible (S) node has at least one excited (E) neighbor, it turns active with a probability depending on the number of its excited neighbors *n*(*E*) and a parameter *p,* which is termed the transmission probability *p*(*S*→*E*) = 1 - (1 - *p*)*^n^*^(*E*)^. This variant of the SER may be called *transmission SER*. The special feature of this model is that, even if a node receives more than one input, and becomes excited in the standard SER, there can be transmission failures in the transmission SER for *P <* 1, and a node can stay susceptible, instead of becoming active. For low values of *p,* the network activity may die out, because of too many transmission failures. For *P* = 1, the model reduces to the deterministic SER with a threshold of one input required to transit from S to E. After a node has been E, it turns refractory (R) in the next time step and is susceptible (S) again in the timestep after that.

In Fig. 6, we show the dynamical behavior of the model when *p* = 1, which is the deterministic SER. Two nodes are excited at the initial time *t* = 0. Activity spreads through the network, and after 10 timesteps, each of the *N* = 192 has fired exactly once (gray crosses, sum E in bottom panel of Fig. 6), and the activity ceases, with all nodes in the fixed point of an S state (green crosses in bottom panel in Fig. 6, #E(t), number of nodes in state E at time t). Thus, for the deterministic setting, the network operates in a through-conduction mode.

**Fig. 6:**
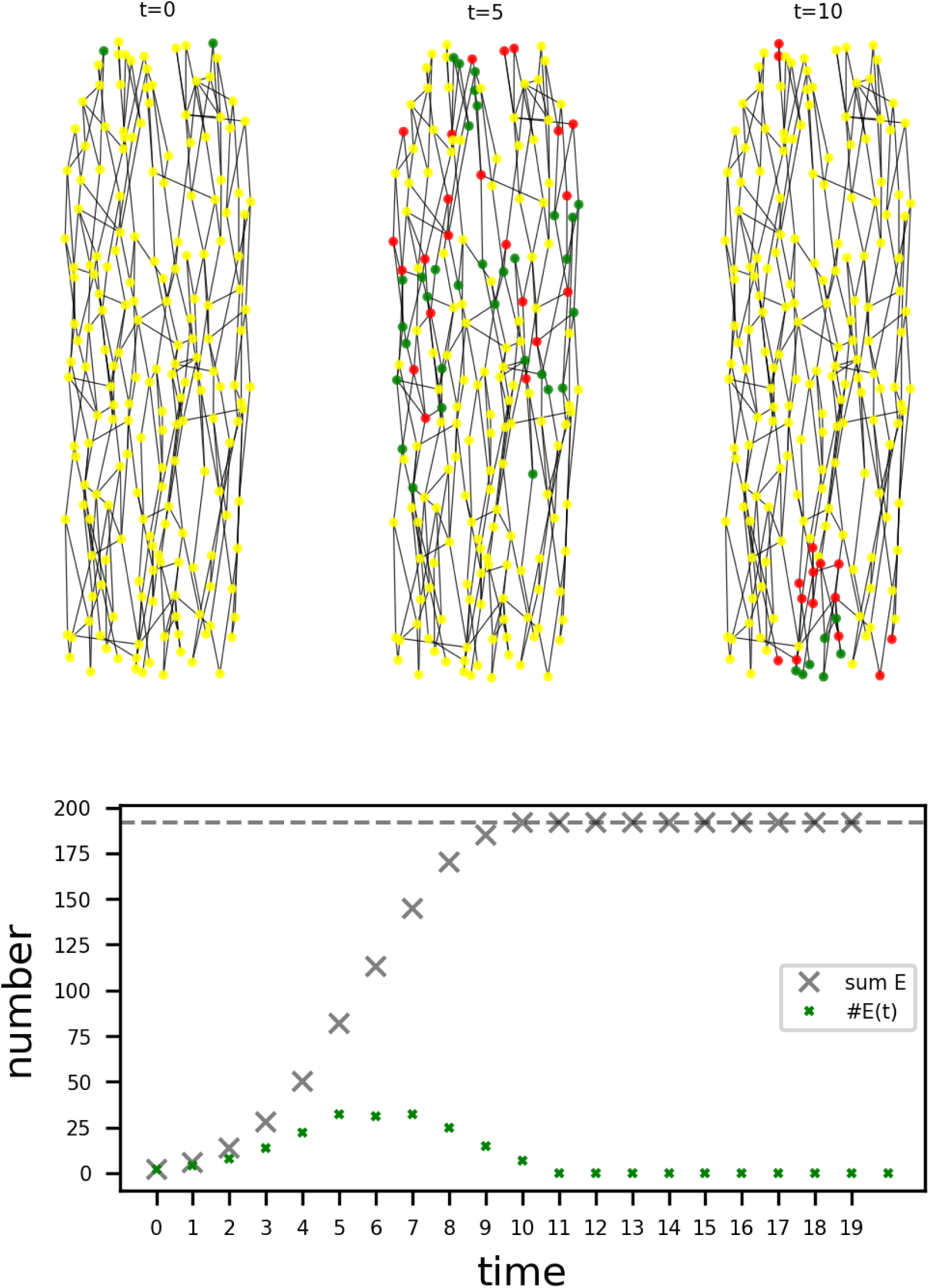
Spread of activity in the spatially embedded network for transmission probability *p* = 10. **Top**: network dynamics at time *t* = 0,5 and io. Color scheme for node states: S- susceptible, E- excited, R- refractory. Activity starts by exciting two nodes at *t* = 0, then traverses through the network. **Bottom:** number of active nodes at time *t*, denoted #E(t) and size of the set of neurons that have fired at least once, denoted sum E. After 10 timesteps, all nodes have fired exactly once (sum E in bottom panel reaches *N* = 192, the total number of neurons in the network, and the sum of# E(t) over all times also reaches *N*), and the network turns silent again. We interpret this as a through conduction mode.

Another activity mode is shown in Fig. 7, where *p* = 0.8, and therefore, transmission failures are expected. In contrast to the dynamics shown in Fig. 6, the network does not turn silent after 10 timesteps. Instead, around 30-50 neurons remain to be active, and the activity persists (bottom panel of Fig. 7, #E(t)). Not all neurons have fired (gray crosses, sum E in bottom panel of Fig. 7), but some neurons have already fired more than once, as the sum of active neurons over time exceeds *N*.

**Fig. 7:**
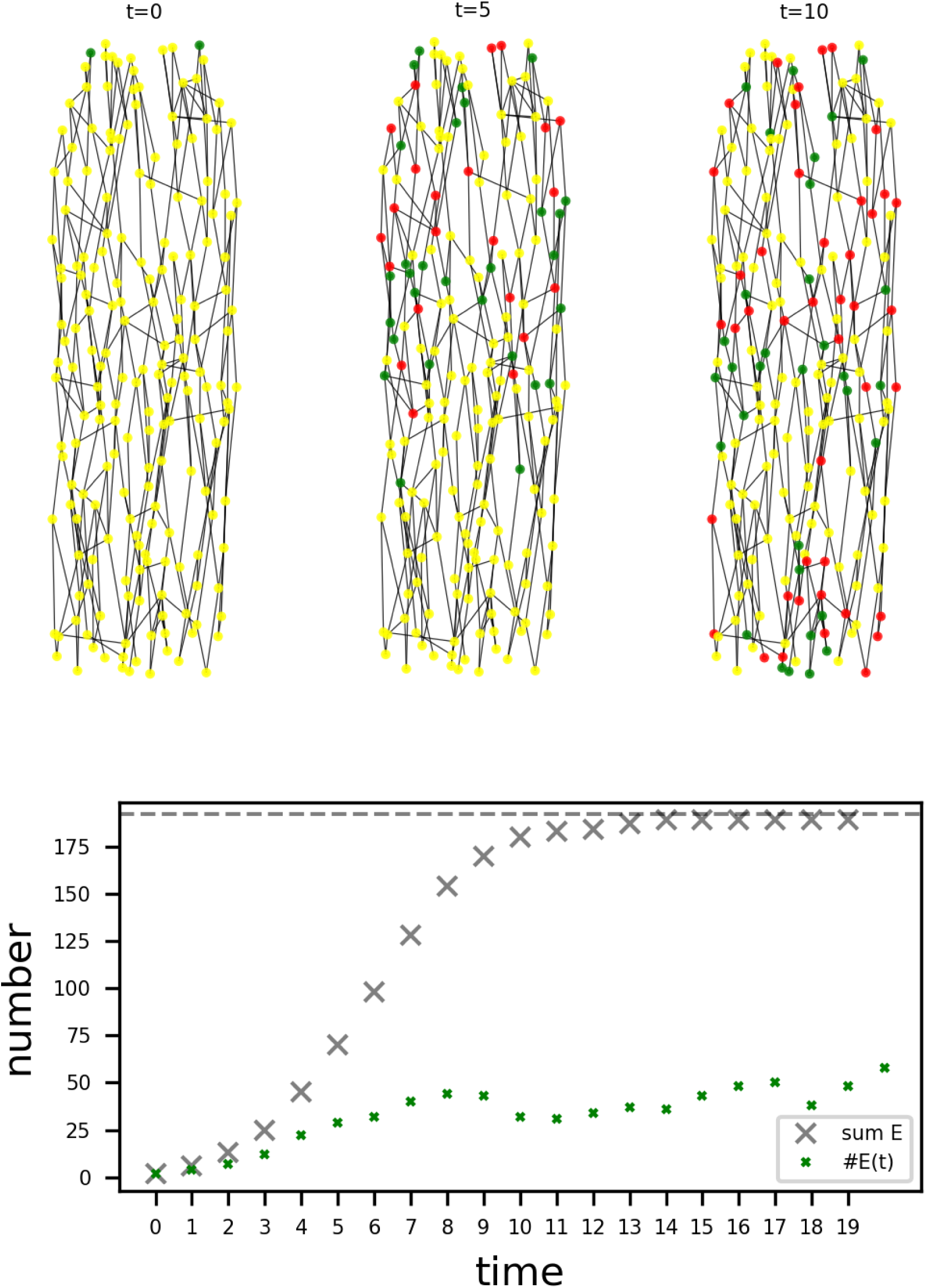
Spread of activity in the spatially embedded network for transmission probability *p* = 0.8. Color code and layout similar to Fig. 6. Note that one node active at *t* = 0 fires again at *t* = 5 (top left), and not all nodes have fired until *t* = 19.

We note that the specific spatial embedding of our connectome model is not directly relevant for the SER model studied here, because there are no temporal delays. However, the topology of the network, as reflected in the specific neighbors of each node, is relevant, and therefore, the generative model indirectly controls the spread of activity in our network.

These observations demonstrate that, already for a simple excitable model ignoring spatiotemporal delays, the network can exhibit at least two qualitatively different modes of activity, one similar to through-conduction also observed in jellyfish neuron models (6) and likely serving information transmission from the peduncle to the hypostome and vice versa, and the other mode similar to persistent spontaneous activity, which were dubbed ‘spontaneous electrical low-frequency oscillations’ (SELFOs) in RP1 and are thought to have a modulatory role in *Hydra*, not generating any observable behavioral output (21–23).

In the next section, we use a more biophysically detailed neuromorphic neuron model to study two modes of activity in our network, where one mode is related to activity propagation, whereas another mode is related to the complex behavioral pattern of somersaulting.

### 2.4 Constructing a functional network model of the body column

Two essential neural activity patterns are known from N3. For the first pattern, activity spreads from the peduncle towards the hypostome or vice versa. For the second pattern, activity spreading from the peduncle to the hypostome drastically increases its firing rate before the foot is detaching and the somersault behavior sequence is initiated. Somersaulting is a complex locomotor behavior in the form of a swaying movement of the whole body column, leading to an attachment of the polyp to an adjacent position (13).

These two patterns (spreading activity and somersaulting) constitute another major aim of this work: Studying to which extent these two activity patterns can be generated by the proposed theoretical connectome and what this implies for neuromorphic hardware realizations. To tackle these questions, we model the somata of the connectome via deterministic neuronal oscillators. To account for a delayed signal transmission due to the distances between the somata, we also model the neurites as an interconnected chain of neuronal oscillators as well. The oscillator model that we deploy is based on (17) and is essentially a Morris-Lecar model in combination with an RC-circuit. The Morris-Lecar model serves to calculate the neural activity in terms of membrane potential, while the RC-circuit provides the calcium-concentration that is qualitatively comparable to fluorescence traces obtained via calcium-imaging measurements.

#### 2.4.1. Modeling the spreading of N3 activity

The Morris-Lecar model used for the spreading activity is described by

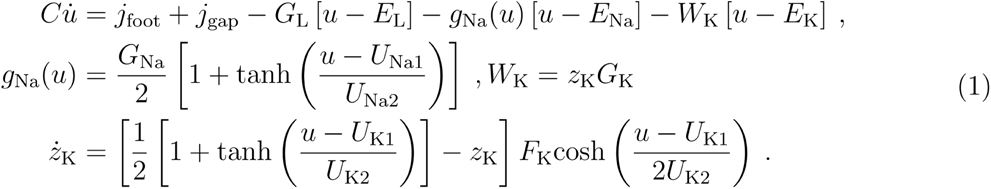

It is computationally less complex than the Hodgkin-Huxley model, but still biologically reasonable and is moreover well interpretable as an electrical circuit, since it naturally comes with an equivalent circuit. Note that we have adopted the representation of the potassium channel as memristor from (24) and have included a sodium channel instead of a calcium-channel to account for the typical ion channel composition of neurons.

The RC-circuit is governed by

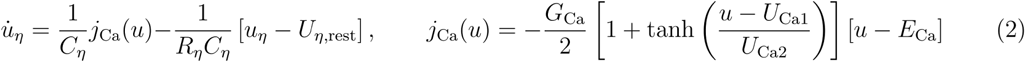

and essentially takes the membrane potential of the Morris-Lecar model as an input signal to generate the calcium concentration. The time constant of the RC-circuit is taken from (17) and is based on the GCaMP6s-data of (25). The resting calcium concentration is chosen as 50 nM following (17) and (26), stating that typical values of neurons range from 50 nM to 100 nM. In combination with the Morris-Lecar model, the circuit representation of the RC-circuit can be found in Fig. 8.

**Fig. 8:**
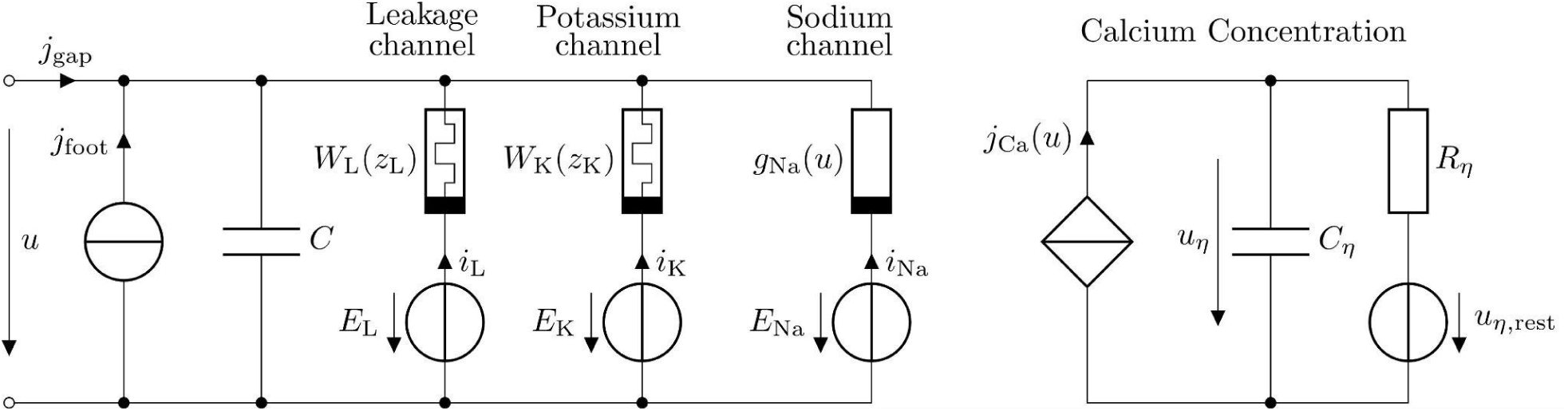
Circuit diagram for Morris-Lecar model (left) and RC-circuit (right).

We create a functional N3 network by placing the described neuronal oscillator at every soma location of the connectome (Fig. 4 and 5), and an interconnected chain of these oscillators as neurites between each pair of somata connected due to the connectome. Here, the interconnections are realized by constant resistors that model the damping of the spatial signal transmission. As couplings between the different somata, we assume gap junctions, because they are known to be largely present in N3 (4). We model these couplings via gap junctions with resistors as well. This is because gap junctions can be interpreted as permanently open ion channels, which are represented by resistors in conductance-based neuron models such as the Morris-Lecar model. For reasons of simplicity, we use the same conductance value *G_e_\* for both gap junction resistors and resistors between the oscillators that form the neurites.

Via circuit simulations of the resulting electrical circuit, this allows us to verify the network’s ability to generate the activity spreading pattern. For this purpose, we excite three somata located near the peduncle by an external, constant current signal. This current accounts for our assumption that the spreading starts from N3 neurons near the foot. Corresponding simulation results can be observed from Fig. 9 and 10. In Fig. 9, we see the membrane potentials of all 192 somata, where low indices correspond to locations near the peduncle and high indices are from somata near the hypostome. Activity starts near the peduncle where the exciting current is applied and then spreads out until it reaches the somata closest to the hypostome. The propagation takes approximately 28 ms. With a body column length of 1.44 mm that constitutes the traveling distance, the conduction velocity is 5.2 cm/s. This fits well with the observed transmission speed of cm/s reported by (21).

**Fig. 9:**
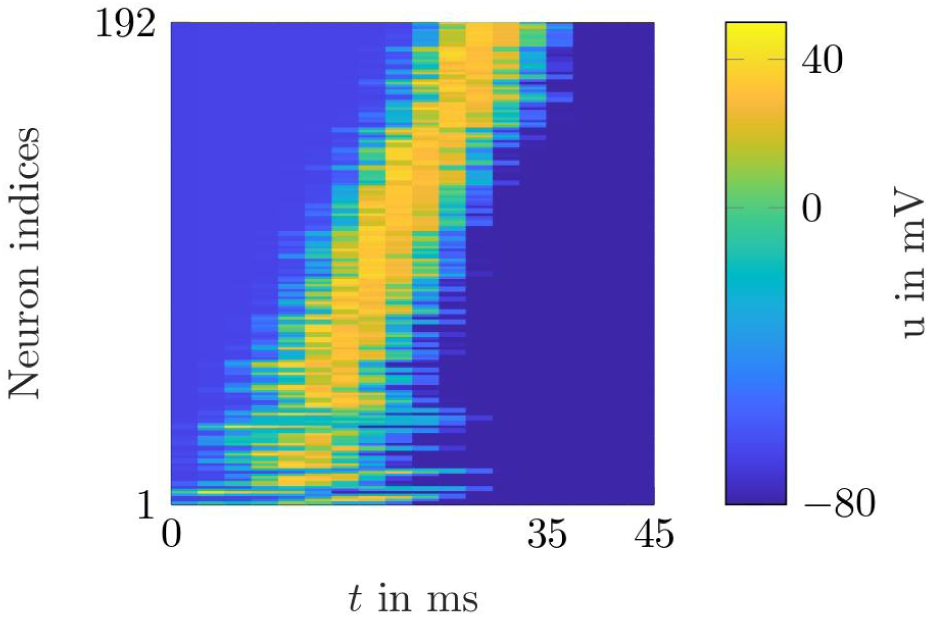
Neural activity in terms of membrane potential spreading from a region close to the peduncle (low neuron indices) to a region near the hypostome (high neuron indices).

**Fig. 10:**
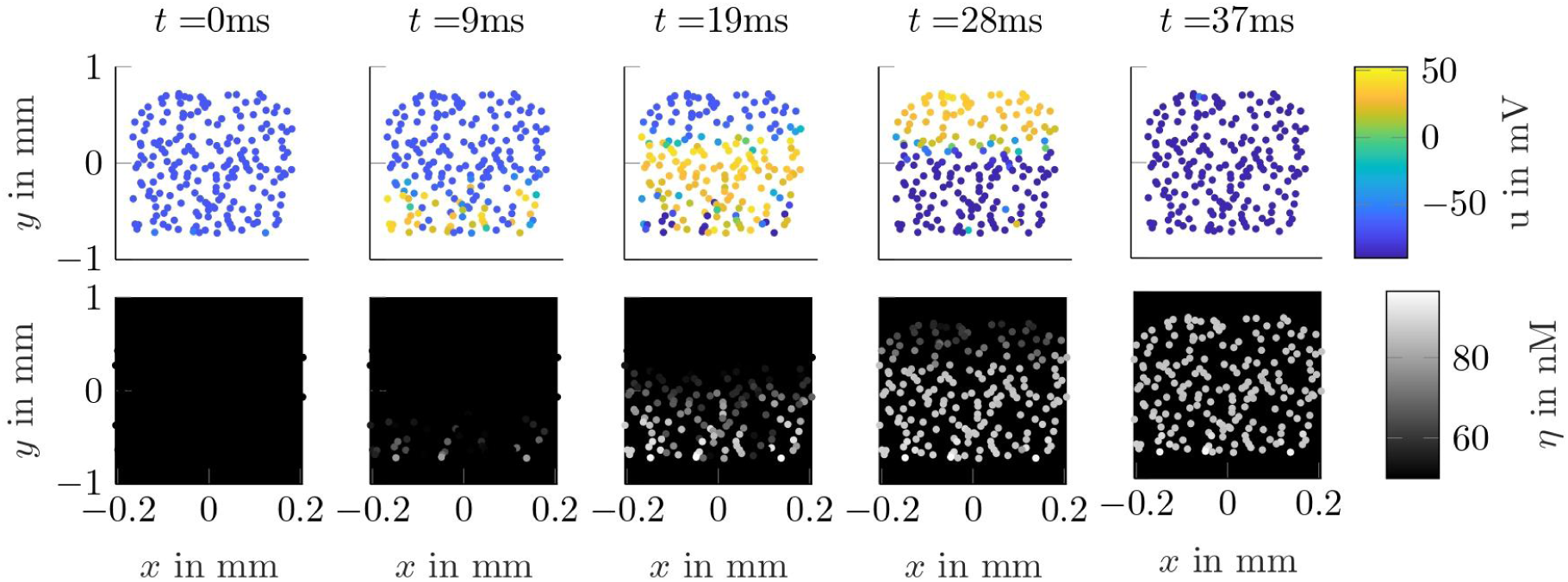
Simulation results for the neural activity spreading across the population N3. Spatial placement as well as dimensions are chosen according to the theoretical connectome. Top: Membrane potentials of the soma. Bottom: Calcium concentrations of the somata.

In Fig. 10, we see the actual network with the somata of the N3 neurons positioned on a 2D grid representing the body column of *Hydra*, like in Fig. 4 and 5. Activity is visualized for five different time points of the simulation, and is shown in terms of both membrane potential and calcium concentration. The membrane potential activity in the top of Fig. 10 clearly lets us identify the currently active somata due to the fast response and decay time of this type of signal. In contrast to this, the calcium-concentration activity in the bottom of Fig. 10 is much slower with respect to its decay time, making it harder to infer which somata are currently active. However, as calcium-imaging is the only widely used measurement method to visualize neuronal activity in *Hydra*, this enables us to generate soma activity that can be compared to measurement data and is also more suitable for representing larger time scales with limited temporal resolution. The latter aspect is especially relevant for neuronal activity patterns related to behavior, which often takes several seconds up to minutes. In the context of this work, such a behavior is the somersaulting motion, which is preceded by an increase in the firing rates of N3 neurons.

#### 2.4.2. Modeling the increase in firing rates of N3 neurons preceding a somersaulting motion

In (13), the spike frequency of the RP1 network has been observed to drastically increase within a period of 5 to 10 minutes and to reach its peak right before somersaulting is initiated by a contraction burst. Characteristic of this frequency behavior is that the increase is weak for most of the time, but extremely strong shortly before the contraction burst. Our aim is now to include this dynamic frequency increase into our functional network model of the body column. As suggested by (13), the frequency modulation is related to the neuropeptide Hym-248. Hym-248 is synthesized by RP1 neurons and belongs to the family of GLWamide peptides. In a recent study, it was shown that Hym-248 is sufficient for somersaulting (13). A detailed mechanism for the increase of firing rates before a somersault is, however, still unknown. It is speculated that Hym-248 mediates a positive feedback loop in RP1, which leads to autostimulation of RP1 (13). For this reason, we hypothesize that a potential mechanism of Hym-248 regulating the frequency behavior is to increase the excitability of the neurons. In this sense, when the concentration of Hym-248 rises due to RP1 activity, the excitability increases and as a result the spike frequencies rises. We implement the dynamic and variable excitability by introducing a dynamic leakage channel. The leakage channel is typically assumed to be constant and mainly determines the force for returning to the resting potential. This directly affects the duration of the depolarization phase and hence the spike frequency. As the leakage channel moreover is a representation of all channels and effects that are not explicitly modeled, it is a well-suited choice to include the influence of Hym-248. To this end, we replace the constant conductance of the leakage channel by a memristor with a linear switching behavior:

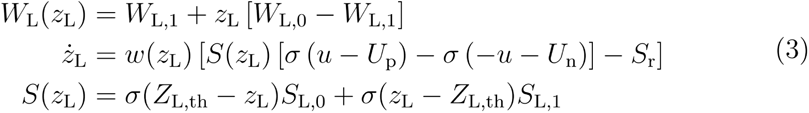

Here, *σ*(·) is the heaviside function and *w*(*z_L_*) ensures the memristor state is at least 0 and at most 1. The described memristor decreases its conductance every time a spike is generated. This in turn increases the excitability and ultimately leads to a rising firing frequency. Moreover, to account for the rapid frequency increase before a beginning contraction burst, we have included a piecewise-linear function for the switching speed of the memristor.

We construct the functional N3 network the same way as before, with the neuronal oscillator being modified with the leakage memristor. The network is again excited by a constant current signal applied to three neurons close to the foot region. This yields the simulation results shown in Fig. 11. In Fig. 11 (a) we see the neuronal activity of all somata in terms of calcium concentration. Note that some spikes are larger than others. These are the ones being directly excited with the current signal. A distinct frequency increase can be inferred from a time point of approximately 320 s. The actual firing frequencies of the sum signal of the calcium concentrations are shown in Fig. 11 (b). Here, a slow increase is present until 300 s, after which a rapid increase takes place. Minimal and maximal observed frequencies are 0.1 Hz and 0.45 Hz, which fits the reports of (13). The dynamics of the frequency change are directly and most probably linearly related to the decreasing conductance of the memristors, see Fig. 11 (c). Note that we have applied a negative current stimulus after 380 s that represents the CB network becoming active, thus terminating the somersault.

**Fig. 11:**
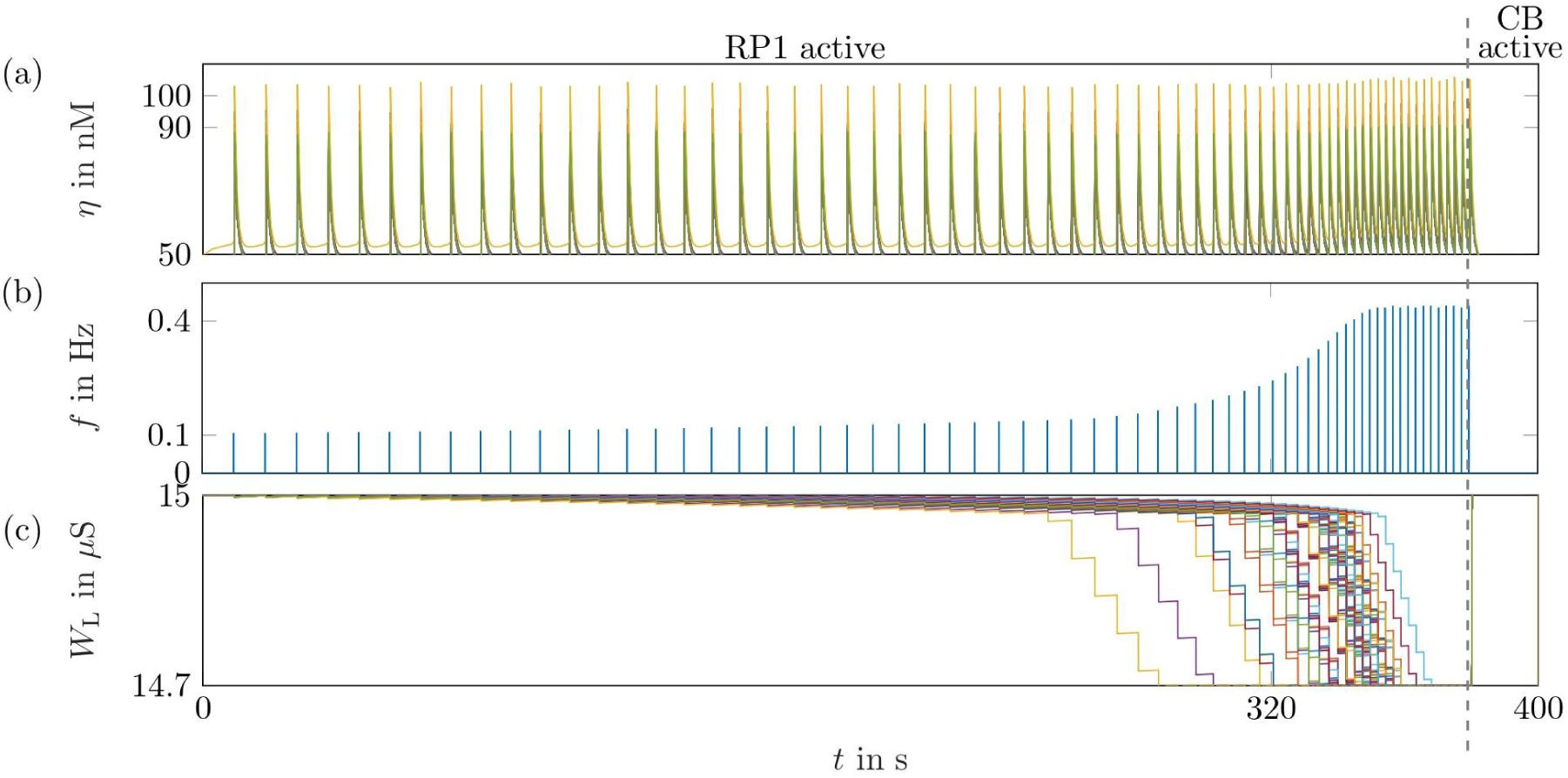
Simulation results for the increasing firing rate of N3 right before the somersaulting motion. (a) Calcium concentrations of all soma. (b) Firing rate of the sum signal of all soma in terms of calcium concentration. (c) Conductance values of the leakage memristors of all soma.

In the present work, we developed a functional network model for the body column neuronal connectivity of *Hydra*, which, despite its seemingly simple organization, has been previously shown to be involved in a variety of activity modes and behaviors. Starting with a number of key experimental observations on the position and orientation of neurons in this network, we constructed a connectome model that best aligned with the structural findings and used two different models for the activity of nodes in this connectome to investigate the resulting network activity patterns. We found that different activity modes underlying specific behaviors in *Hydra* could be obtained by small parameter variations in the models, suggesting that a combination of overall connectome topology and local dynamical node tuning may be responsible for the rich behavioral repertoire associated with the body column network. Our approach, combining empirical and computational observations, has a number of specific implications and limitations, as discussed in the following sections.

This study constructed and validated a functional network model of a body column neuronal network of the cnidarian polyp *Hydra*. Taking a geometric perspective, we used a generative network model to distill the network topology in the form of a representative model connectome. We initially focussed on the body column neuronal network N3, which is also identified with the network RP1 (9,13). Our modeling approach reveals several emergent features: generating neuron positions using a modified PD sampling approach (Figs. 2 and 3), we designed connectivity rules that only assume the maximal number of connections per neuron, the preferred direction of connections and the maximal connection length as input. We showed that this information is sufficient to reproduce key experimental data, including the distribution of connection lengths and connection angles (Fig. 4). The derived mean number of primary neurites per neuron (between 2 and 3, Fig. 4, degree/ primary neurites) is similar to that observed for the RP1 network by (9). However, their neuronal densities in the body column are higher than ours (more than 1300 vs. 321 mm ^-2^). However, in their Fig. S3, densities are plotted per population and RP1 has a mean density of around 500 mm^-2^, which is closer to our value. Moreover, their orientation data showed no preferred orientation like in our model. The difference in density and orientation can likely be attributed to various factors. First, the genetic construct used in that study was not specific only to N3/RP1, which could result in higher neuron numbers and hence higher neuron densities (cf. Fig. 4A in (9), were no elongated neurites and no preferred neurite orientation is visible). The non-specificity of the staining in general makes it hard to compare their data to our recordings. Second, their animals appear to be in a contracted state, which influences their body shape and therefore likely also cellular density and orientation measurements.

Next, we focused on studying dynamics on the generated network model. Using the transmission SER model, we showed that already this simple model generated two different activity patterns, one similar to through conductance (Fig. 6), which may be used for signaling along the top-to-bottom axis, and the other one qualitatively similar to spontaneous activity (Fig. 7), both of which are observed in real *Hydra* (22). Interestingly, a model assuming imperfect synaptic transmission, as given by the transmission SER with activation probability smaller than 1, gave rise to persisting activity in the model N3 network (Fig. 7), whereas a model assuming perfect synaptic transmission (Fig. 6) gave rise to through-conductance of activity, in which every neuron only fired once.

We then applied a neuromorphic circuit approach to study activity propagation in our model connectome. We found that the spatially embedded network could give rise to two different activity modes, one linked to through-conductance of the system, and the other one linked to the onset of the somersaulting behavior. The exact mechanism of frequency modulation via Hym-248 is not clear. While we focus on the binding of Hym-248 to RP1 neuron receptors reducing firing thresholds, it is potentially also possible that Hym-248 modulates gap junction activity. There is no direct evidence for gap junctions being modulated by neuropeptides in Hydra, however, cf. (27), where it was shown that an ion channel is directly activated by neuropeptides in the *Hydra* nervous system. Gap junctions are important for synchronization and have also been reported to regulate network-wide oscillations and their frequency, see (28). In this sense, a weakening of gap junctions via Hym-248 could lead to an asynchronous spiking behavior that results in more detected spikes for a sum signal of the RP1 network.

How could our connectome model be improved? Accepting that in the absence of precise synaptic markers and without electron microscopy imaging for *Hydra* (see, however, (14–16)), a ground-truth validation of the connectome is not possible at the moment, there seem to be three main directions of improvement. First, a more realistic three-dimensional spatial model could be designed. In doing so, the complex neurite geometry, which we have chosen to ignore in this paper, could be incorporated, and a comparison with the simplified two-dimensional model could be performed. Second, a comparison between different node dynamical systems could reveal which node model best reproduces measured calcium traces. In this direction, neuron models of the Hodgkin-Huxley (HH) type could be designed, which show how different ion channels in the neuronal membrane give rise to spiking activity. This detailed biophysical modeling approach is, however, only beneficial when intracellular electrophysiology data become available for *Hydra* neurons. Specifically, our approach is in principle already able to generate results comparable to calcium activity measurements, and without electrophysiologically characterized neurons, HH-type models do not increase the biological accuracy, but only computational complexity. The reason for this is that the presence or absence and the voltage dependence of ion channels cannot be validated without intracellular electrophysiology, and therefore, their precise mathematical form for *Hydra* neurons remains elusive. Third, neuro-muscular coupling, similar to (20), could be modeled to achieve a more complete model of *Hydra* behavior.

In this work, we have exclusively focused on the comparatively simple body column network N3. The complexity of the neuronal networks is higher in the hypostome and in the foot. Especially in the peduncle, it is assumed that cross-talk between the nerve nets N1 and N3 occurs (11), thus coordinating *Hydra* behavior. Likewise, in the hypostome, it was recently shown that interaction between networks N3, N4, and N6 is crucial for the complex feeding behavior of *Hydra* (11). Recent work also showed that N1 and N3 each control different aspects of feeding behavior in underfed *Hydra* (12). It is an open task to model the structure and dynamics of different subnetworks in order to understand their interplay, and ultimately their relation to *Hydra* behavior. Another exciting direction for future work is to study the dynamic maintenance of *Hydras* nerve net. In *Hydra*, neurons in the body column are constantly replaced by neuronal precursor cells derived from abundant interstitial stem cells (29–33). As a result, every neuron only has a finite life span of at most several weeks after it has entered the network and become functional. Our modeling approach considered the *Hydra* nerve net as a static arrangement of cells, which in light of the constant neuronal turnover does not seem to be reflecting biological reality. It seems more appropriate to consider the network we have presented in this work as a snapshot at one particular point in time. Future work could therefore focus on modeling the cell cycle and the migration of neuronal precursor cells to better understand the dynamic maintenance of the nerve net and the preservation of its network topology over time.

Finally, the complex ontogenesis of *Hydra* could be studied using generative modeling. Re-aggregating *Hydra* were studied in (34), where it was shown that the whole *Hydra* nerve net can reassemble from small cellular aggregates, leading to global network synchronization via growing assemblies of neurons. Combining our generative modeling approach with models for activity-dependent network plasticity and growth rules thus seems like another fruitful direction of research.

An interesting perspective on *Hydra’s* nerve net was recently provided by (16). In this work, it was shown that nerve nets are formed by bundles of parallel overlapping neurites, which are linked by circuit-specific gap junctions (16). It was also suggested that the body column neuronal network in *Hydra* could maintain its structure and grow by a process termed lateral addition (16), in which newborn neurons first grow their neurites in a direction essentially parallel to the long axis of the animal and then attach to the existing network when their neurites overlap with those of already integrated neurons. For our modeling approach, it offers the possibility to study outgrowth models, in which connections of different strengths are formed whenever two neurites overlap, and the more overlap there is, the stronger the connection will be. Our algorithm to place somata on a two-dimensional domain could be used in models of that type. The crucial step will be to devise biologically plausible growth rules for neurites that take into account chemoattractants and neuronal growth cones. Neurites in *Hydra* could also be so-called anastomosed structures, and that its nerve net therefore forms a syncytium, with no clear differentiation between neurites and neuronal somata. This was recently shown for the nervous system of ctenophora (35), which are a sister group to all animals with nervous systems.

The fact that *Hydra* uses neuropeptides to control somersaulting (13) might point to a role of non-synaptic information transmission in its nervous system. Extrasynaptic information transmission was recently shown to be crucial for understanding activity propagation in the nematode *C. elegans* in the form of a ‘wireless connectome’ (36,37). In light of these findings, one might be tempted to question the relevance of any connectome model studying physical connections between neurons as the basis of neuronal communication in *Hydra*. Still, the contribution of our paper can be seen to consist of the functional verification of basic mechanisms of activity propagation in *Hydra*, which requires the connectome for the functional model. To understand in detail how neuropeptides act on single neurons and during network dynamics in conjunction with a physical connectome requires more data on N3 activity and its modulation.

Our work offers a systematic construction of the structure of a subnetwork of *Hydra’s* nervous system. By taking the view that the ectodermal body column network in *Hydra* is essentially two-dimensional, we designed a generative network model that is in agreement with measured structural quantities and supports two different activity modes, each presumably controlling different types of behavior in *Hydra*.

## 4. Acknowledgments

WB would like to thank Kayson Fakhar, Fatemeh Hadaeghi and Mariia Popova for helpful discussions. Funded by the Deutsche Forschungsgemeinschaft (DFG, German Research Foundation) - Project-ID 434434223 - SFB 1461. Research in the laboratory of T.C.G.B. is supported in part by grants from the German Research Foundation (Deutsche Forschungsgemeinschaft, DFG), the CRC 1182 ‘Origin and Function of Metaorganisms’ (to T.C.G.B.), and the CRC 1461 ‘Neurotronics: Bio-Inspired Information Pathways’ (Project-ID 434434223 - SFB 1461) to T.C.G.B. and A.K. T.C.G.B. appreciates support from the Canadian Institute for Advanced Research.

## 5. Methods

### 5.1. Wave digital simulation

Circuit simulations are carried out by translating the mathematical model constituting an electrical circuit into a wave digital model. This is based on the wave digital concept introduced by (38). A wave digital model is obtained by port-wisely decomposing the electrical circuit and then translating each device as well as their interconnection structure into the wave digital domain using the bijective transformation

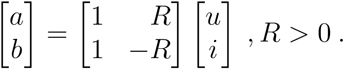

Following this procedure for the electrical circuit shown in Fig. 8 and described by equations (1), (2) and (3), this yields the vector-valued wave digital model shown in Fig. 12. For more details on the model derivation, the interested reader is referred to (17,39,40). The model is programmed with Matlab.

**Fig. 12:**
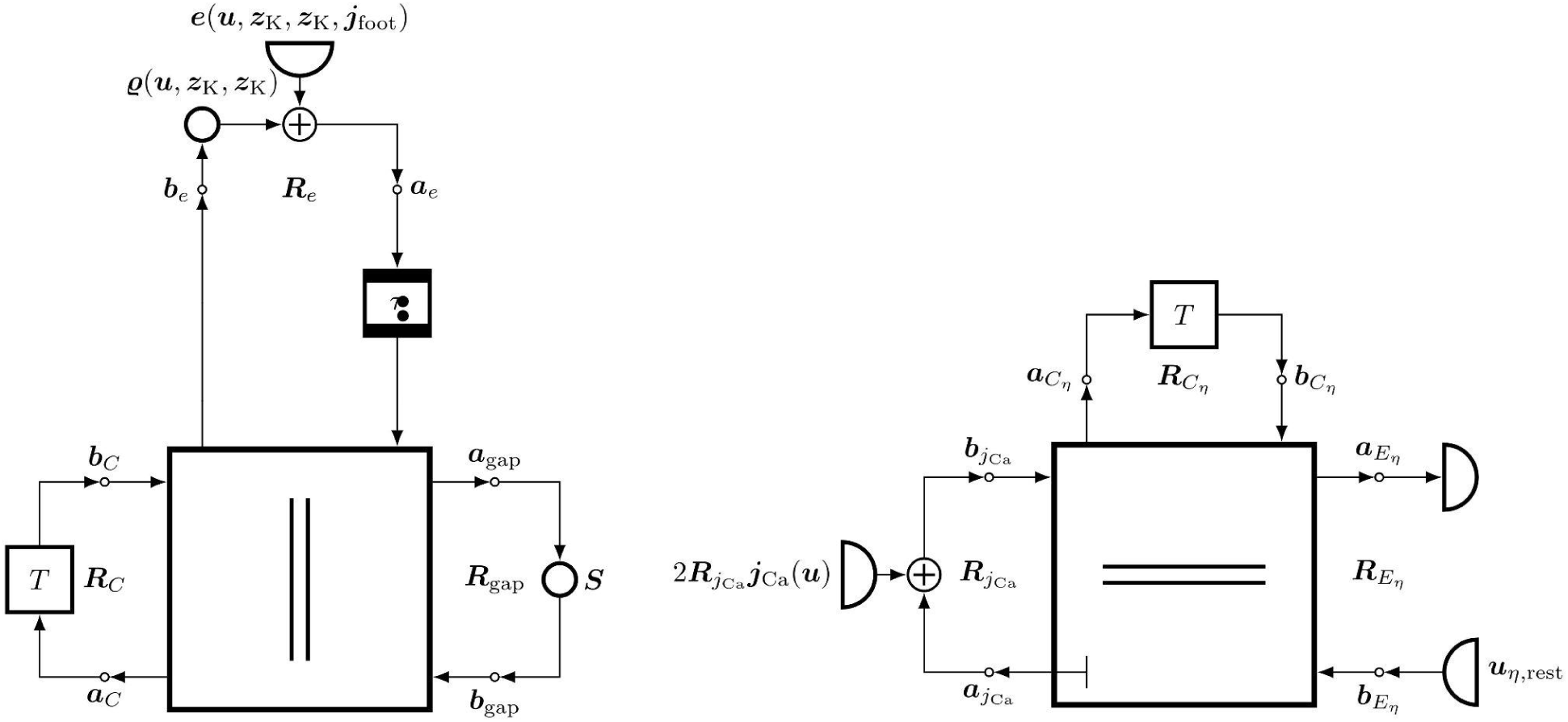
Vector-valued wave digital model.

### 5.2. Coupling structure of functional network model via adjacency and incidence matrices

The coordinates of the somata as well the adjacency matrix describing their coupling structure are derived from the theoretical connectome. The adjacency matrix serves as a starting point for a graph-theoretical description of the coupling structure, while the coordinates are used to determine the distances between the nodes of the graph. To account for spatially extended neurons and hence signal transmission delays, we create a new graph. In particular, we represent each edge *e* of the original graph by *k*(*e*) nodes and edges of the new graph that account for the neurite length between two somata. *k*(*e*) is determined by dividing the distance between the two considered somata by a constant factor chosen to 50 and rounding the result to the next integer. Based on the new graph, we derive an incidence matrix, since this is required for the wave digital model of the coupling structure, see (40).

### 5.3 Determining spike frequencies

Spike frequencies of the simulated circuit model are obtained by first calculating the sum signal of the calcium concentrations of all somata. Here, the sum signal allows us to generate results comparable to (13). The sum signal is then processed with the *findpeaks* function of Matlab. The distances of the detected peaks are used to calculate the frequency.

### 5.4. *Hydra* maintenance and transgenic animals

*Hydra vulgaris* AEP was maintained in accordance to standard procedures in standard *Hydra* culture medium (CaCl2 0.042g/L; MgSO4x7H20 0.081g/L; NaHCO3 0.042g/L, K2CO3 0.011g/L in dH2O) (Ref: https://doi.org/10.1038/s41596-019-0173-3). Animals were fed three times per week with *Artemia salina* and maintained at 18 C with a 12h/12h light cycle.

Transgenic animals expressing GCaMP6S under the promoter of the gene t12874aep, specific to N3 neurons, were derived in another study (11) and maintained as wild type animals. Animals were starved for at least two days before an experiment.

### 5.5. Immunohistochemistry against GFP and Imaging

To visualize the neuronal network N3 and analyze the structure and properties of the network, transgenic animals expressing GCaMP6S under the promoter for N3 were fixed and stained with an antibody against GFP. First, animals were relaxed with 2% urethan and fixed for 2h at RT or overnight at 4°C in Zamboni (Morphisto, cat#12773). Here the crucial step is the treatment with urethan, as it determines the relaxation stage of the animals, which influences the downstream analysis. Second, animals were washed in 1x PBS with 0.1% Tween (PBST) followed by an incubation in 1xPBS with 0.5% TritonX100 and a 1h blocking step in PBST with 1% BSA. Third, the primary antibody chicken-anti-GFP (Biozol, cat# GFP-1010, 1:1000 dilution) was added and incubated overnight at 4°C. Fourth, animals were washed in PBST with 1% BSA before the secondary antibody was added. The secondary antibody goat anti-chicken Alexa Fluor 488 (Invitrogen, cat# A11039, 1:1000 dilution) was added and incubated for2h at RT. After washing in PBST with 0.5% Tween and 1% BSA, animals were incubated for 5 min in TO-PRO-3 Iodide (642/661) (Invitrogen, cat#T3605, 1:1000 dilution). And last, animals were mounted in moviol with DAPCO on glass slides and stored at 4°C till imaging. Slides were imaged by using the Axio Vert. A1 (Zeiss) with Colibri 7 as a light source (Zeiss) equipped with the fluorescence filter 38 HE (Zeiss). The 5x, 10x, and 20x Plan Apo objectives together with the Axiocam 705 mono (Zeiss) were used to take the images.

### 5.6. Analysis of neuronal population N3

To determine the density, number of neurites, the angle of the connections, length of connections, and soma-soma connections we processed the images with ImagJ (Fiji; doi:10_i_1038/nmeth_J_2019). All images were oriented and aligned in a first step and further adjusted (Brightness/Contrast, Window/Level) to enhance the visibility of all neurites and somata. Cells were counted using the CellCounter plugin. Tracing of neurites was done using the SNT package (doi:10.1038/s41592-021-01105-7).

Angles are measured with a sign (see Fig. S1): neurites/connections pointing upwards, i.e. in the quadrant I and II of a cartesian coordinate system, are assigned positive angles 0° ≤ *β* ≤ 180°; angles pointing downwards (in the quadrants III and IV of a cartesian coordinate system) are assigned negative angles -180° ≤ *β* ≤ 0°. In the model, two angles are recorded for each made connection, one as given with the first neuron at the origin of a cartesian coordinate system, the other one as given with the second neuron at the origin. These angles will differ by 180° and therefore, the distribution of angle values in the model will be symmetric around 0°.

**Fig. S1:**
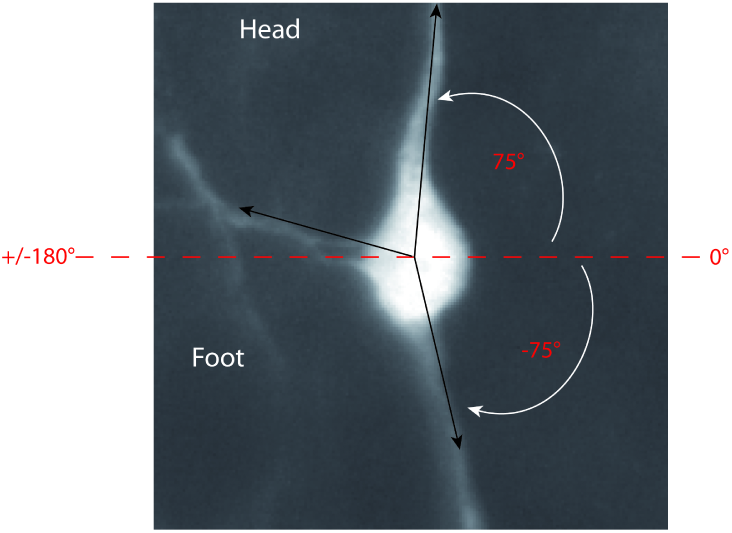
Sign conventions for angles. Neurites (connections) pointing upwards are assigned positive angle values, connections pointing downwards are assigned negative values. Note that while this depiction shows oriented neurites, actual measurements were performed as shown in Fig. S2.

**Fig. S2:**
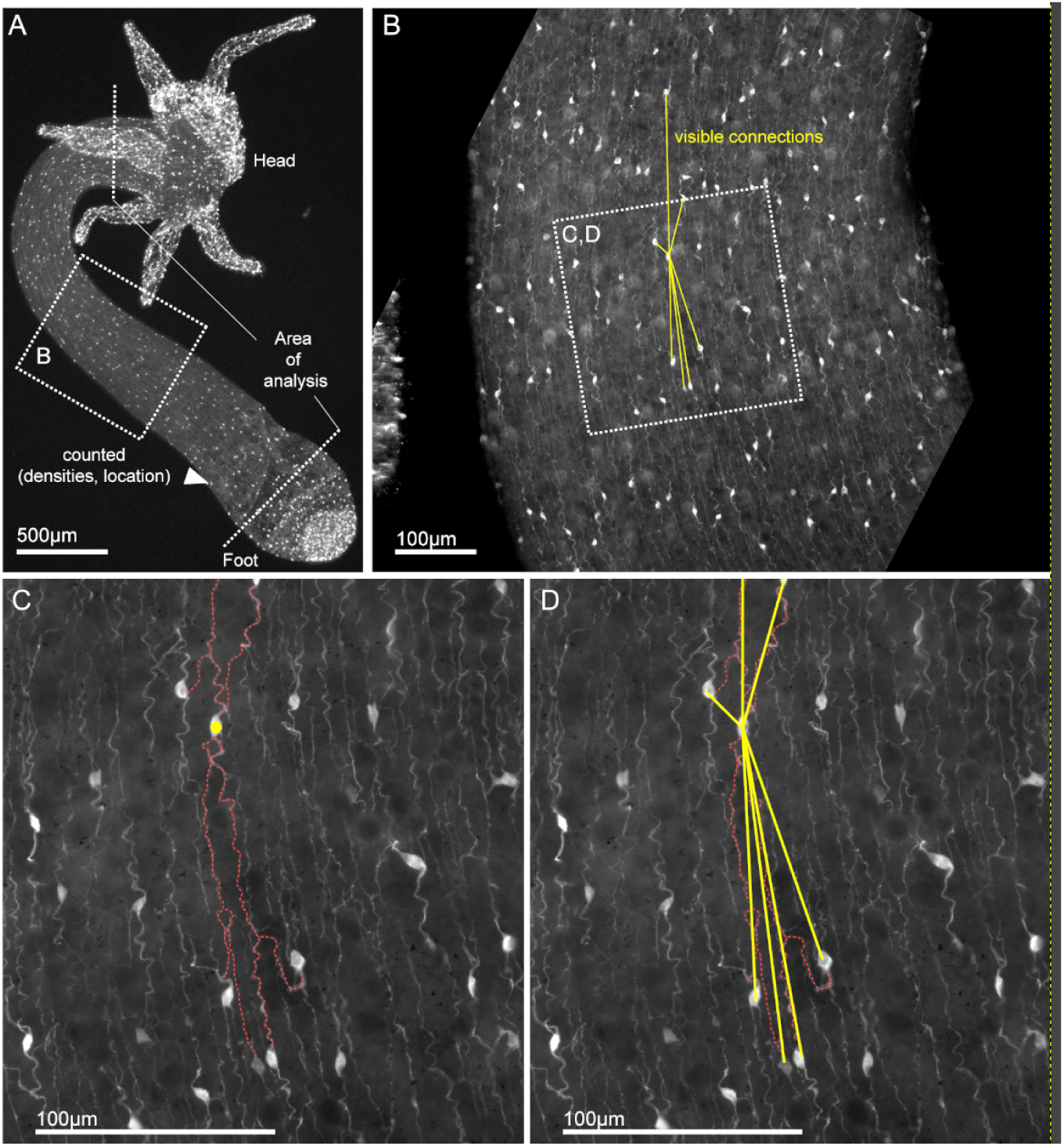
Experimental setup. (A) *Hydra* polyp with head, body column and foot. Section of the body column shown in (B) is marked with a dotted rectangle. Density and location of neurons was counted in the entire body. (B) Neurites were traced in only a section of the body column for multiple polyps. Connections between two neurons were deemed existent whenever two neurites overlapped, either directly or via branching. The direct Euclidean distance of connected neurons was then measured, and compared to the model connectome output. (C) Neurite tracing for one neuron, shown in yellow. (D) Measurement of connection distances (yellow lines) for one neuron. The neuron depicted here has 7 connections.

## Appendix

### A. Parameter tables for circuit models

**Table 1.**
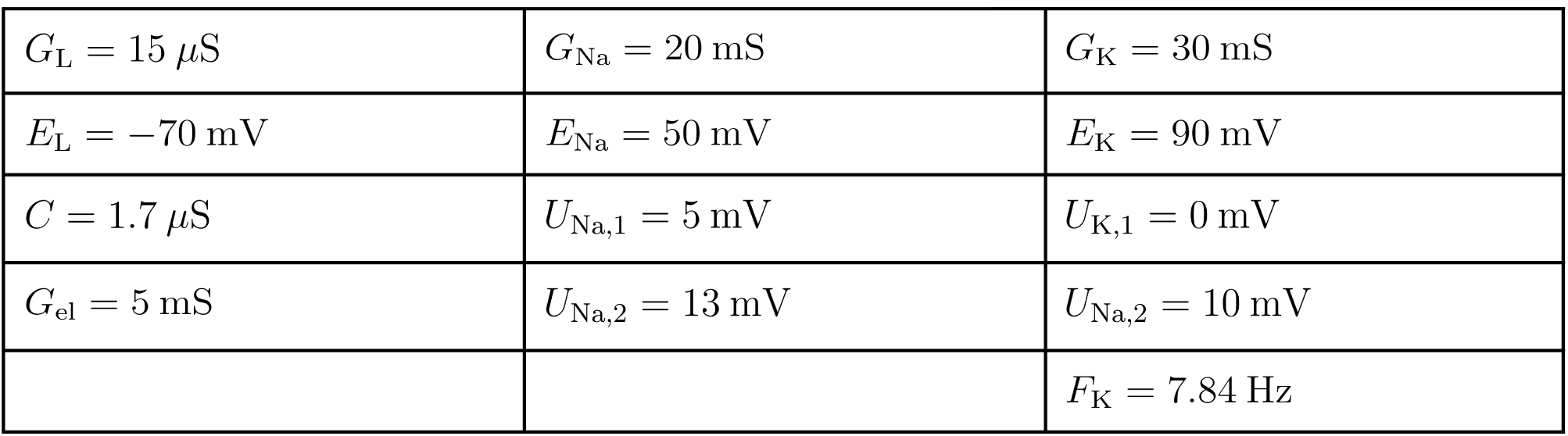
Circuit parameters for the Morris-Lecar model.

|  |  |  |
| --- | --- | --- |
| $G_L = 15 \mu\text{S}$ | $G_{\text{Na}} = 20 \text{ mS}$ | $G_K = 30 \text{ mS}$ |
| $E_L = -70 \text{ mV}$ | $E_{\text{Na}} = 50 \text{ mV}$ | $E_K = 90 \text{ mV}$ |
| $C = 1.7 \mu\text{S}$ | $U_{\text{Na},1} = 5 \text{ mV}$ | $U_{K,1} = 0 \text{ mV}$ |
| $G_{\text{el}} = 5 \text{ mS}$ | $U_{\text{Na},2} = 13 \text{ mV}$ | $U_{\text{Na},2} = 10 \text{ mV}$ |
| | | $F_K = 7.84 \text{ Hz}$ |

**Table 2.** Circuit parameters for the memristor model of the leakage channel.

|  |  |  |  |
| --- | --- | --- | --- |
| $W_{L,0} = 14.7 \mu\text{S}$ | $W_{L,1} = 15 \mu\text{S}$ | $U_p = 10 \text{ mV}$ | $U_p = -100 \text{ mV}$ |
| $S_{L,0} = 0.83 \mu\text{Hz}$ | $S_{L,0} = 16.67 \mu\text{Hz}$ | $S_r = 0.83 \text{ mHz}$ | |

**Table 3.**
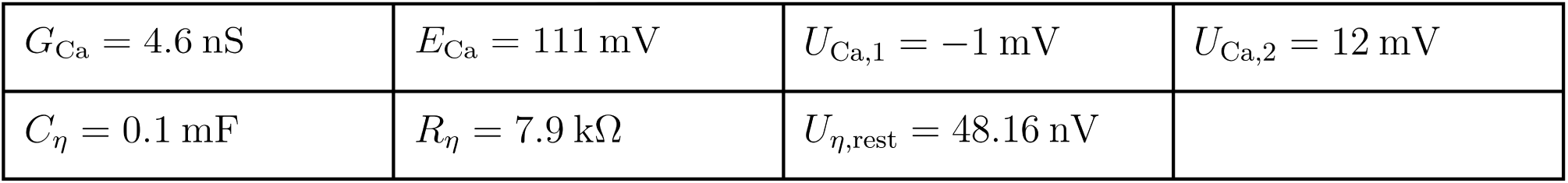
RC circuit parameters.

## Bibliography

1. Kandel ER, Schwartz JH, Jessell TM, Siegelbaum SA, Hudspeth AJ. Principles of Neural Science. 5. edition. New York: McGraw-Hill Education Ltd; 2012. 1760 p.

2. White JG, Southgate E, Thomson JN, Brenner S. The structure of the nervous system of the nematode Caenorhabditis elegans. Philos Trans R Soc Lond B Biol Sci. 1986 Nov 12;314(1165):1-340.

3. Siebert S, Farrell JA, Cazet JF, Abeykoon Y, Primack AS, Schnitzler CE, et al. Stem cell differentiation trajectories in Hydra resolved at single-cell resolution. Science. 2019 Jul 26;365(6451):eaav9314.

4. Klimovich A, Giacomello S, Björklund Â, Faure L, Kaucka M, Giez C, et al. Prototypical pacemaker neurons interact with the resident microbiota. Proc Natl Acad Sci USA. 2020 Jul 28;117(30):17854–63.

5. Bosch TCG, Klimovich A, Domazet-Lošo T, Gründer S, Holstein TW, Jékely G, et al. Back to the Basics: Cnidarians Start to Fire. Trends Neurosci. 2017 Feb 1;40(2):92–105.

6. Pallasdies F, Goedeke S, Braun W, Memmesheimer RM. From single neurons to behavior in the jellyfish Aurelia aurita. Calabrese RL, Satterlie R, Kanso E, editors. eLife. 2019 Dec 23;8:e50084.

7. Weissbourd B, Momose T, Nair A, Kennedy A, Hunt B, Anderson DJ. A genetically tractable jellyfish model for systems and evolutionary neuroscience. Cell. 2021 Nov 24;184(24):5854–5868.e20.

8. Bielecki J, Dam Nielsen SK, Nachman G, Garm A. Associative learning in the box jellyfish Tripedalia cystophora. Curr Biol. 2023 Oct;33(19):4150–4159.e5.

9. Dupre C, Yuste R. Non-overlapping Neural Networks in Hydra vulgaris. Curr Biol. 2017 Apr 24;27(8):1085–97.

10. Campbell RD. Elimination of Hydra interstitial and nerve cells by means of colchicine. J Cell Sci. 1976 Jun 1;21(1):1–13.

11. Giez C, Pinkle D, Giencke Y, Wittlieb J, Herbst E, Spratte T, et al. Multiple neuronal populations control the eating behavior in Hydra and are responsive to microbial signals. Curr Biol. 2023 Dec;33(24):5288–5303.e6.

12. Giez C, Noack C, Sakib E, Hofacker LM, Repnik U, Bramkamp M, et al. Satiety controls behavior in Hydra through an interplay of pre-enteric and central nervous system-like neuron populations. Cell Rep [Internet]. 2024 Jun 25 [cited 2024 May 31];43(6). Available from: https://www.cell.com/cell-reports/abstract/S2211-1247(24)00538-2

13. Yamamoto W, Yuste R. Peptide-driven control of somersaulting in Hydra vulgaris. Curr Biol. 2023 May 22;33(10):1893–1905.e4.

14. Kinnamon JC, Westfall JA. Types of neurons and synaptic connections at hypostome-tentacle junctions in Hydra. J Morphol. 1982;173(1):119–28.

15. Westfall JA, Epp LG. Scanning electron microscopy of neurons isolated from the pedal disk and body column of Hydra. Tissue Cell. 1985 Jan 1;17(2):161–70.

16. Keramidioti A, Schneid S, Busse C, Laue CC von, Bertulat B, Salvenmoser W, et al. A new look at the architecture and dynamics of the Hydra nerve net. eLife [Internet]. 2023 Jun 20 [cited 2023 Nov20];12. Available from: https://elifesciences.org/reviewed-preprints/87330

17. Jenderny S, Ochs K. Wave digital model of calcium-imaging-based neuronal activity of mice. Int J Numer Model Electron Netw Devices Fields. 2023;36(2):e3053.

18. Bridson R. Fast Poisson disk sampling in arbitrary dimensions. In: ACM SIGGRAPH 2007 sketches [Internet]. New York, NY, USA: Association for Computing Machinery; 2007 [cited 2023 Nov 21]. p. 22-es. (SIGGRAPH ’07). Available from: 10.1145/1278780.1278807

19. Goulas A, Betzel RF, Hilgetag CC. Spatiotemporal ontogeny of brain wiring. Sci Adv. 2019 Jun;5(6):eaav9694.

20. Wang H, Swore J, Sharma S, Szymanski JR, Yuste R, Daniel TL, et al. A complete biomechanical model of Hydra contractile behaviors, from neural drive to muscle to movement. Proc Natl Acad Sci. 2023 Mar 14;120(11):e2210439120.

21. Hanson A. On being a Hydra with, and without, a nervous system: what do neurons add? Anim Cogn [Internet]. 2023 Aug 4 [cited 2023 Aug 10]; Available from: 10.1007/s10071-023-01816-8

22. Hanson A. Spontaneous electrical low-frequency oscillations: a possible role in Hydra and all living systems. Philos Trans R Soc B Biol Sci. 2021 Jan 25;376(1820):20190763.

23. Passano LM, McCullough CB. THE LIGHT RESPONSE AND THE RHYTHMIC POTENTIALS OF HYDRA. Proc Natl Acad Sci USA. 1962 Aug;48(8):1376–82.

24. Chua L, Sbitnev V, Kim H. Hodgkin-huxley axon is made of memristors. Int J Bifurc Chaos. 2012 Mar;22(03):1230011.

25. Chen TW, Wardill TJ, Sun Y, Pulver SR, Renninger SL, Baohan A, et al. Ultrasensitive fluorescent proteins for imaging neuronal activity. Nature. 2013 Jul;499(7458):295-300.

26. Grienberger C, Konnerth A. Imaging Calcium in Neurons. Neuron. 2012 Mar 8;73(5):862–85.

27. Gründer S, Assmann M. Peptide-gated ion channels and the simple nervous system of Hydra. J Exp Biol. 2015 Feb 15;218(Pt 4):551–61.

28. Pernelle G, Nicola W, Clopath C. Gap junction plasticity as a mechanism to regulate network-wide oscillations. PLOS Comput Biol. 2018 Mar 12;14(3):e1006025.

29. David CN, Campbell RD. Cell cycle kinetics and development of Hydra attenuata. I. Epithelial cells. J Cell Sci. 1972 Sep;11(2):557–68.

30. David CN, Hager G. Formation of a primitive nervous system: nerve cell differentiation in the polyp hydra. Perspect Dev Neurobiol. 1994;2(2):135–40.

31. Teragawa CK, Bode HR. Migrating interstitial cells differentiate into neurons in hydra. Dev Biol. 1995 Oct;171(2):286–93.

32. Hager G, David CN. Pattern of differentiated nerve cells in hydra is determined by precursor migration. Dev Camb Engl. 1997 Jan;124(2):569–76.

33. Bosch TCG, Anton-Erxleben F, Hemmrich G, Khalturin K. The Hydra polyp: nothing but an active stem cell community. Dev Growth Differ. 2010 Jan;52(1):15–25.

34. Lovas JR, Yuste R. Ensemble synchronization in the reassembly of Hydra’s nervous system. CurrBiol CB. 2021 Sep 13;31(17):3784–3796.e3.

35. Burkhardt P, Colgren J, Medhus A, Digel L, Naumann B, Soto-Angel JJ, et al. Syncytial nerve net in a ctenophore adds insights on the evolution of nervous systems. Science. 2023 Apr 21;380(6642):293-7.

36. Randi F, Sharma AK, Dvali S, Leifer AM. Neural signal propagation atlas of Caenorhabditis elegans. Nature. 2023 Nov;623(7986):406-14.

37. Ripoll-Sánchez L, Watteyne J, Sun H, Fernandez R, Taylor SR, Weinreb A, et al. The neuropeptidergic connectome of C. elegans. Neuron. 2023 Nov 15;111(22):3570–3589.e5.

38. Fettweis A. Wave digital filters: Theory and practice. Proc IEEE. 1986 Feb;74(2):270–327.

39. Ochs K, Michaelis D, Jenderny S. An Optimized Morris-Lecar Neuron Model Using Wave Digital Principles. In: 2018 IEEE 61st International Midwest Symposium on Circuits and Systems (MWSCAS) [Internet]. 2018 [cited 2023 Dec 15]. p. 61-4. Available from: https://ieeexplore.ieee.org/document/8623905

40. Ochs K, Michaelis D, Roggendorf J. Circuit Synthesis and Electrical Interpretation of Synchronization in the Kuramoto Model. In: 2019 30th Irish Signals and Systems Conference (ISSC) [Internet]. 2019 [cited 2023 Dec 15]. p. 1-5. Available from: https://ieeexplore.ieee.org/document/8904942

